# Update on the reproduction and interpretation of DIANA fMRI

**DOI:** 10.1101/2025.08.31.673005

**Authors:** Jae-Youn Keum, Semi Park, Heejung Chun, Daehong Kim, Jang-Yeon Park

## Abstract

Three years ago, our group reported direct imaging of neuronal activity (DIANA) with high spatiotemporal resolution, but its reproducibility and signal origin remain controversial. Here, we report the results of our reproduction experiments of DIANA fMRI performed to date using forelimb electrical stimulation at various magnetic field strengths in anesthetized mice, along with the characteristics of DIANA signal, called the pseudo-steady state (PSS). Theoretical analysis and Bloch simulations demonstrated that the spatial location and temporal phase of PSS oscillations are primarily determined by frequency-offset, and that their spatiotemporal superposition can generate peak signals in specific regions at specific timing that closely resemble DIANA signals. These findings suggest that if PSS oscillations are the primary source of DIANA signals, it may be premature to interpret them as neuronal responses to sensory stimulation. Further studies are needed to clarify the relationship between DIANA signals and brain activation.

## Introduction

Three years ago, our group reported a novel functional magnetic resonance imaging (fMRI) method called Direct Imaging of Neuronal Activity (DIANA), which allows direct detection of neuronal activity on millisecond timescales using a two-dimensional (2D) gradient-echo line-scan MR imaging strategy (*1*). In this original study, we showed that DIANA fMRI with 5 ms temporal resolution could capture neuronal activity and its propagation in thalamocortical (TC) pathway of the somatosensory network during whisker-pad electrical stimulation in anesthetized mice at 9.4 T. The first DIANA response was observed sequentially in the thalamus and primary somatosensory barrel field (S1BF) at ∼10 ms and ∼25 ms, respectively, consistent with previous electrophysiological studies (*2–4*).

DIANA fMRI has attracted significant attention as a groundbreaking attempt to noninvasively capture neuronal activity on millisecond timescales *in vivo*, a long-standing goal of neuroscientists. However, in the midst of a journey toward that goal, difficulty in replicating the DIANA signal has been reported (*5–9*). Several groups trying to reproduce the DIANA signal have failed in animal (*5*, *6*) and human studies (*7–9*). These results raised doubts regarding the reproducibility of DIANA fMRI and led to considerable criticism of the initial publication. Despite such criticism and loss of hope, some groups have reported the observation of DIANA signal. Meng et al. have observed the DIANA signal in the somatosensory cortex of awake mice at 9.4 T (*10*) and our group has also reported detection of DIANA signals in somatosensory pathways at 7 T (*11–13*) and 11.7 T (*14–17*), as well as in the visual pathways of mice at 9.4 T (*18*).

On the other hand, the DIANA signal was also criticized for possibly being an artifact signal due to nonideal aspects of the pulse sequence. Phi Van et al. reported that the delay time assigned to trigger stimulation in a DIANA pulse sequence can disrupt the steady state of the spin system and cause undesired peak signals that are not related to neuronal activation (*19*). They made the important discovery that even small trigger delays (e.g., 12 μs) in the line-scan pulse sequence of DIANA fMRI can produce unwanted peak signals, the peak timing of which is somewhat delayed with decreasing spin-spin relaxation time (T_2_) and static magnetic field (B_0_) inhomogeneity. However, this report could not fully explain the characteristics of the DIANA signal, particularly the temporal correlation between DIANA and electrophysiological signals in specific regions. For example, in our original study (*1*), the peak DIANA response was observed earlier in the thalamus than in S1BF, where T_2_ and B_0_ inhomogeneity are higher due to a greater blood-oxygenation-level-dependent (BOLD) contribution during brain activation.

In this study, we report important recent investigations into the properties of the DIANA signal, including the results of our reproduction experiments of DIANA fMRI performed to date using forelimb electrical stimulation at various magnetic field strengths (e.g., 11.7 T and 7 T) in anesthetized mice. Importantly, we report an physical mechanism that contributes significantly to DIANA signal generation, called a pseudo-steady state (PSS) of magnetization due to RF spoiling (*20–22*). The periodic oscillations induced by the PSS of magnetization (hereafter referred to as PSS oscillations) appear to depend spatially and temporally on frequency-offsets, providing a plausible explanation for the DIANA signal patterns observed in the gradient-echo line-scan acquisition scheme. However, at this stage, it remains premature to conclude that the frequency-offset dependence of PSS oscillations is primarily attributed to neuronal activity, as further investigations are required.

## Results

### DIANA responses at 11.7 T with trigger delay and RF spoiling

To check whether *in vivo* DIANA signals can be reproduced at different magnetic field strengths, we performed DIANA fMRI on an 11.7 T animal scanner (BioSpec 117/16 USR, Bruker BioSpin) using six adult C57BL/6J medetomidine-anesthetized mice (*14*) (Fig. 1A). A 2D fast low-angle shot (FLASH) line-scan imaging sequence was used, as in the original study (*1*). Sufficient dummy scans (8 s, 1600 TRs) were performed before the main scan to achieve steady-state magnetization (*23*). Both radiofrequency (RF) spoiling (*24–26*) and gradient spoiling (*27*) were also used to suppress the effects of residual transverse magnetization (**M***_xy_*). A trigger delay of 25 μs was introduced, as in the original study (*1*). We delivered electrical stimulation (strength, 0.5 mA; duration, 1 ms; frequency, 5 Hz) to the right forelimb of the mouse with an interstimulus interval of 200 ms, consisting of 50 ms pre-stimulation, 1 ms stimulation, and 149 ms post-stimulation (Fig. 1B). Stimuli were applied 64 times corresponding to the number of phase-encoding lines. A single 0.8 mm thick oblique slice covering both the thalamus and forelimb primary somatosensory cortex (S1FL) was acquired to observe responses in the thalamus and S1FL simultaneously with scan parameters as follows (Fig. 1C): repetition time (TR) = 5 ms (temporal resolution), echo time (TE) = 2 ms, flip angle (FA) = 4°, field of view (FOV) = 25.6 × 12.8 mm^2^, and spatial resolution = 0.2×0.2×0.8 mm^3^. 40 trials per mouse were acquired and used for analysis. To minimize neural adaptation, a 90-second rest period was provided every 5 trials.

**Fig. 1.**
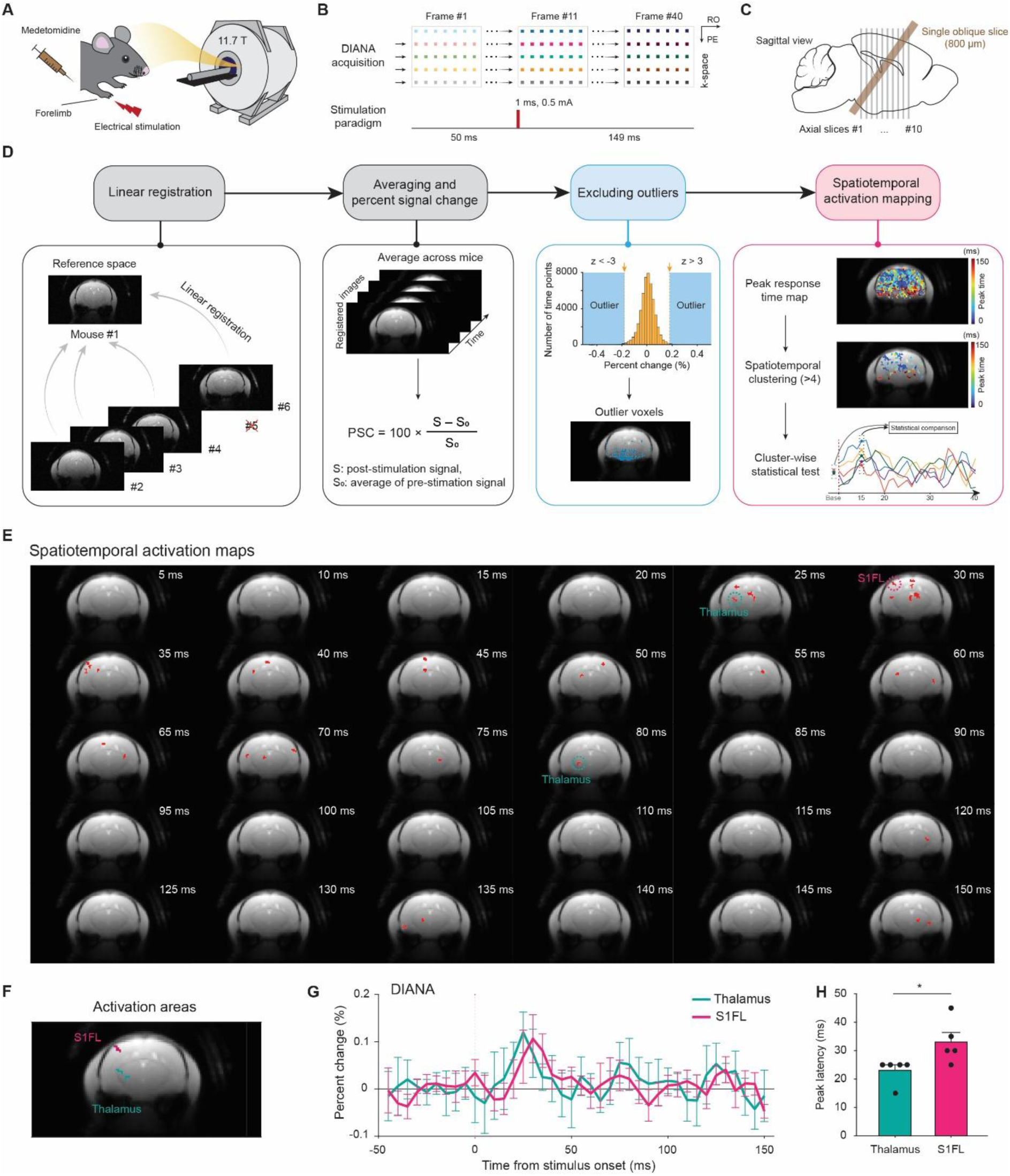
Spatiotemporal activation mapping in DIANA fMRI. **(A to B)** Schematics of the DIANA fMRI experiment (A), 2D line-scan acquisition and stimulation paradigm (B) to capture DIANA responses to electrical right forelimb stimulation in medetomidine-anesthetized mice at 11.7 T. **(C)** Location of the set oblique slice in sagittal plane. **(D)** Flowchart of the spatiotemporal activation mapping process: Linear registration of DIANA images of 4 mice (#2 to #6, excluding #5 due to no BOLD response in the thalamus) to the mouse #1 image as template. Average the data across all mice and represent the time series as percent signal change (PSC). Exclude outlier voxels with z > 3 or z < −3 using a histogram of the number of voxels with respect to PSC. Spatiotemporal activation mapping consisting of generation of a peak response time map, spatiotemporal clustering (cluster size > 4), and cluster-wise testing for statistical significance of DIANA responses. **(E)** Spatiotemporal activation maps of DIANA fMRI with 5 ms temporal resolution after stimulus onset. **(F)** Representative DIANA-activated regions within the thalamus (cyan) and S1FL (magenta). **(G to H)** DIANA time series (G) and mean latencies (H) of DIANA response peaks obtained from the DIANA-activated regions within the thalamus and S1FL (*n* = 5 mice, *: *p* < 0.05, one-tailed paired *t*-test). Vertical dotted lines indicate the onset time of electrical forelimb stimulation (H). All data are mean ± SEM.

Prior to DIANA fMRI, scout BOLD fMRI was performed as a reference to identify the location of the thalamus and S1FL, and to ensure reliable neuronal activation in response to sensory stimulation (fig. S1). BOLD activation maps showed highly consistent activation within the contralateral thalamus and contralateral S1FL across the mice, except for the thalamus in mouse #5. Since the oblique slice of mouse #5 appeared to be improperly setup to not include the thalamus, mouse #5 was excluded from further analysis.

To perform group analysis on 5 mice, each DIANA image for mice #2 to #6 was linearly registered to the DIANA image for mouse #1 and then averaged (Fig. 1D, leftmost). First, averaging across all voxels within the brain, we observed a peak signal change of 0.034% at 30 ms after trigger onset (*n* = 5 mice), which appears to be related to the trigger delay (fig. S2).

For DIANA time series analysis, we developed a data analysis method (termed “spatiotemporal activation mapping”) to identify DIANA activation regions in an effective and reliable manner by exploiting the temporal information of DIANA responses, which is a key advantage of DIANA fMRI over conventional fMRI (Fig. 1D). In other words, the peak response time map was created by obtaining the times of peak DIANA responses on a voxel-by-voxel basis from the time series of the group-averaged DIANA image, and clustered voxels above a cluster size of 4 were selected as activation regions. An important rationale for this spatiotemporal activation mapping is the assumption that spatially clustered voxels of a certain size in the peak response time map can be considered a group of functionally identical features sharing the same response time and cannot be generated by chance. As a final step, we statistically compared the DIANA signal at all post-stimulus time points to the pre-stimulus baseline to test the statistical significance of the DIANA response for the mean time series of each cluster. Clusters that survived this statistical test were considered to represent the final spatiotemporal activation areas.

Prior to the spatiotemporal activation mapping, voxels with at least one data point with z score > 3 or < −3 were excluded as outliers, considering the percent signal change (PSC) histogram of all data points in the time series for all voxels. Here, the PSC was defined as a percentage of the post-stimulus measurement relative to the average of the data points in the pre-stimulus period. The outlier voxels were mostly concentrated in the lower part of the brain (Fig. 1D, second from the right), where the temporal signal-to-noise ratio (tSNR), defined as the signal mean divided by signal standard deviation over the 40 frames, was relatively low because a surface coil was used on top of the brain (fig. S2). This is because high PSC values corresponding to z score > 3 or < −3 may be due to low average values in the pre-stimulus period in the low tSNR region. The maximum and minimum values of the tSNR map obtained from the average time series of 200 trials (= 5 mice × 40 trials/mouse) were 1469.4 and 104.17, respectively. The mean tSNR value of the outlier voxels was 371.33 ± 104.04.

As a result, the spatiotemporal distribution of all DIANA activation areas was shown in Fig. 1E, including the somatosensory network. Among all activation areas, the TC pathway from the ventral posterolateral nucleus (VPL) of the thalamus to S1FL was first presented in Fig. 1F to H. In response to the electrical forelimb stimulation, peak DIANA responses were sequentially observed in the VPL and S1FL with peak latencies of 23.0 ± 2.00 ms and 33.0 ± 3.39 ms, respectively (Fig. 1H, *n* = 5 mice, *: *p* < 0.05, paired *t*-test), which was in good agreement with previous electrophysiological studies using mouse forelimb stimulation (*28–35*).

While these results appear to well reproduce DIANA fMRI at 11.7 T, the latencies of peak signals in the VPL and S1FL were in a similar range to the latency of peak signal induced by trigger delay (∼30 ms), observed when averaging across all voxels in the brain. Therefore, as a next step, we investigated whether the DIANA response could persist even after removing the trigger delay and at different magnetic field strengths.

### DIANA responses at 7 T without trigger delay and with RF spoiling

To adjust the trigger delay, we modified the FLASH line-scan sequence source code (ppg file), where the trigger delay is caused by two delays introduced into each of the two trigger functions, located immediately before and after the first TR, respectively. In the original FLASH imaging sequence, these two delays were each set to 3 μs each by default.

To experiment with different trigger delay values, we modified the second delay assigned to the second trigger function located immediately after the first TR (Δ*t* in Fig. 2A). On the other hand, to remove the trigger delay, both delays assigned to the two trigger functions located immediately before and after the first TR were both modified to 0 μs, and the trigger functions previously outside the TR loop (required for line-scan) were also moved inside the TR loop to eliminate any possibility of trigger delay due to the trigger function (Fig. 2C).

**Fig. 2.**
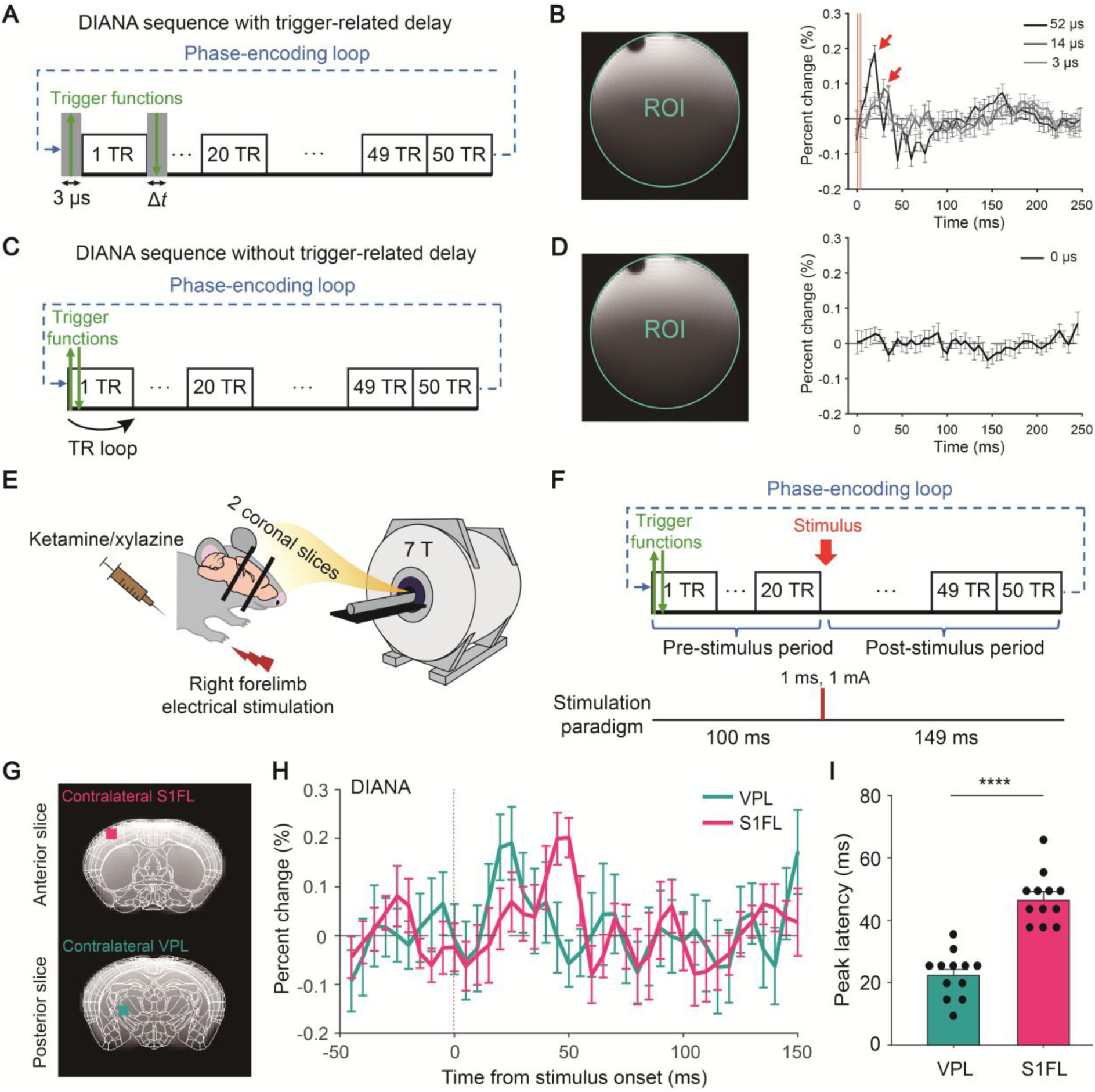
DIANA fMRI in ketamine/xylazine-anesthetized mice at 7 T. **(A)** Schematics of DIANA pulse sequence including trigger function present right before and after the first TR. **(B)** Comparison of DIANA time courses averaged over the entire phantom region with respect to delay assigned to the trigger function right after the first TR (50 trials). **(C)** Schematics of a modified DIANA pulse sequence with the delay removed and the trigger functions moved inside the TR loop. **(D)** DIANA time courses averaged over the entire phantom region obtained with the modified DIANA sequence (50 trials). **(E to F)** Schematics of the DIANA fMRI experiment (E) and stimulation paradigm (F) to capture DIANA responses to electrical right forelimb stimulation in ketamine/xylazine-anesthetized mice at 7 T. **(G)** Square ROIs selected for the VPL (cyan) and S1FL (magenta) after overlaying the Allen Mouse Brain Atlas template. **(H to I)** DIANA time courses (H) and mean latencies (I) of DIANA response peaks obtained from the ROIs within the thalamus and S1FL (*n* = 12 mice, ****: *p* < 0.0001, one-tailed Welch’s *t*-test). Vertical red dotted lines indicate the onset time of electrical forelimb stimulation (H). All data are mean ± SEM except for (B) and (D) which are mean ± SD.

After implementing the modified sequences on a 7 T animal scanner (BioSpec 70/20 USR, Bruker BioSpin), we performed phantom imaging using the following scan parameters: TR = 5 ms (temporal resolution), TE = 2 ms, FA = 4.6°, FOV = 16 × 16 mm^2^, and spatial resolution = 0.22 × 0.22 × 1.0 mm^3^. Other scan parameters were the same as those used in 11.7 T DIANA fMRI introduced in the previous chapter.

First, DIANA signals were acquired using sequences with different trigger delays, and the results were shown in Fig. 2B. When using a saline phantom with a trigger delay of 55 μs (1^st^ delay, 3 μs by default; 2^nd^ delay, 52 μs), a signal change with a peak of ∼0.19 % was observed, and it decreased with decreasing trigger delay, i.e., ∼0.09% and ∼0.05% with a trigger delay of 17 μs (= 3 μs + 14 μs) and 6 μs (= 3 μs + 3 μs), respectively. All peak signal changes were observed around 20 - 30 ms after trigger onset.

Next, to verify that the trigger delay was removed, the modified line-scan imaging sequence was implemented on the same scanner and phantom imaging was performed again with the same scan parameters. As a result of removing the assigned delay and moving the trigger functions inside the TR loop, we confirmed that the signal change due to the trigger delay was well suppressed as shown in Fig. 2D.

Finally, to investigate whether the stimulus-responsive *in vivo* DIANA signals are still observable without trigger delay, DIANA fMRI was performed using 12 adult ketamine/xylazine-anesthetized Cre-inducible diphtheria toxin receptor transgenic mice (iDTR) mice on the same animal scanner (Fig. 2E). We used iDTR mice because we originally planned to observe changes in neural circuits involved in the somatosensory network before and after inducing reactive astrocytes in the VPL using forepaw electrical stimulation (*12*).

We delivered electrical stimulation (strength, 1 mA; duration, 1 ms; frequency, 4 Hz) to the right forelimb of the mouse with an interstimulus interval of 250 ms, consisting of 100 ms pre-stimulation, 1 ms stimulation, and 149 ms post-stimulation, to achieve event-synchronized line-scan acquisition (Fig. 2F). Stimuli were applied 72 times corresponding to the number of phase-encoding lines. Two coronal slices covering the VPL and S1FL, respectively, were imaged with the same scan parameters as those used for phantom imaging. 40 trials per mouse were acquired and used for analysis. To minimize neural adaptation, a 90-second rest period was provided every 5 trials.

Using the sequence with trigger delay removed, we still identified the TC pathway from the VPL to S1FL (Figs. 2G to I), showing that DIANA fMRI at 7 T is reproducible even without trigger delay. In response to the electrical forelimb stimulation, peak DIANA responses were sequentially observed in the VPL and S1FL with peak latencies of 22.5 ± 1.91 ms (∼0.19 %) and 47.9 ± 1.99 ms (∼0.20 %), respectively (*n* = 12 mice, Fig. 2I, ****: *p* < 0.0001, Welch’s *t*-test). It is interesting to note that there was a 10 - 20 ms delay in the DIANA response to forelimb stimulation in both VPL and S1FL compared to whisker pad stimulation originally reported in (*1*), likely due to the longer pathway in the forelimb sensory circuit, and this result is in good agreement with previous electrophysiological studies using mouse forelimb stimulation (*28–35*).

Considering these results together with the observations above at 11.7 T, it seems to suggest that although the previously reported DIANA response may have been influenced to some extent by the trigger delay effect, it cannot be fully explained by the trigger delay effect alone and that other factors may also be involved. In the following sections, we will introduce another important factor that significantly contributes to the DIANA signals, along with the underlying physical mechanism.

### PSS M**_xy_** due to RF spoiling in gradient-echo sequences

In fact, we discovered another important factor mentioned above that significantly influences the DIANA signals while analyzing human DIANA fMRI data. In human DIANA fMRI studies from our group and others (*7*), prominent ∼28 Hz and ∼56 Hz frequency components were observed in the DIANA time series, particularly near the brain periphery. Since these periodic signal components can seriously influence the interpretation of functional signals in the time series, we attempted to elucidate their cause and finally found that they are closely linked to the imperfect steady-state (or PSS as mentioned earlier) signal behavior associated with RF spoiling in gradient-echo sequences (*20–22*). In this section, we first briefly introduce the theory of PSS signal behavior due to RF spoiling, focusing on how periodic signal oscillations are generated.

Conventional gradient-echo sequences utilize RF spoiling in addition to gradient spoiling to more effectively suppress the effects of residual **M***_xy_* (Fig. 3A). In the FLASH sequence, RF spoiling is implemented through quadratic RF phase cycling, which is expressed as *ϕ*_n_ = *n*(*n*−1)*ψ*/2, where *ϕ*_n_ is the phase of the *n*^th^ excitation RF pulse and *ψ* is the phase increment of RF spoiling, which is typically set to 50° or 117° (117° was used in our study). The receiver phase in each TR is set to be equal to *ϕ*_n_. According to previous studies (*20–22*), the quadratic phase cycle of RF spoiling causes the spin system to reach a PSS of **M***_xy_* immediately after RF excitation, satisfying the following relation,

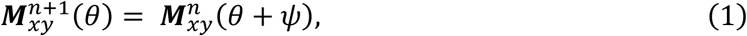

**Fig. 3.**
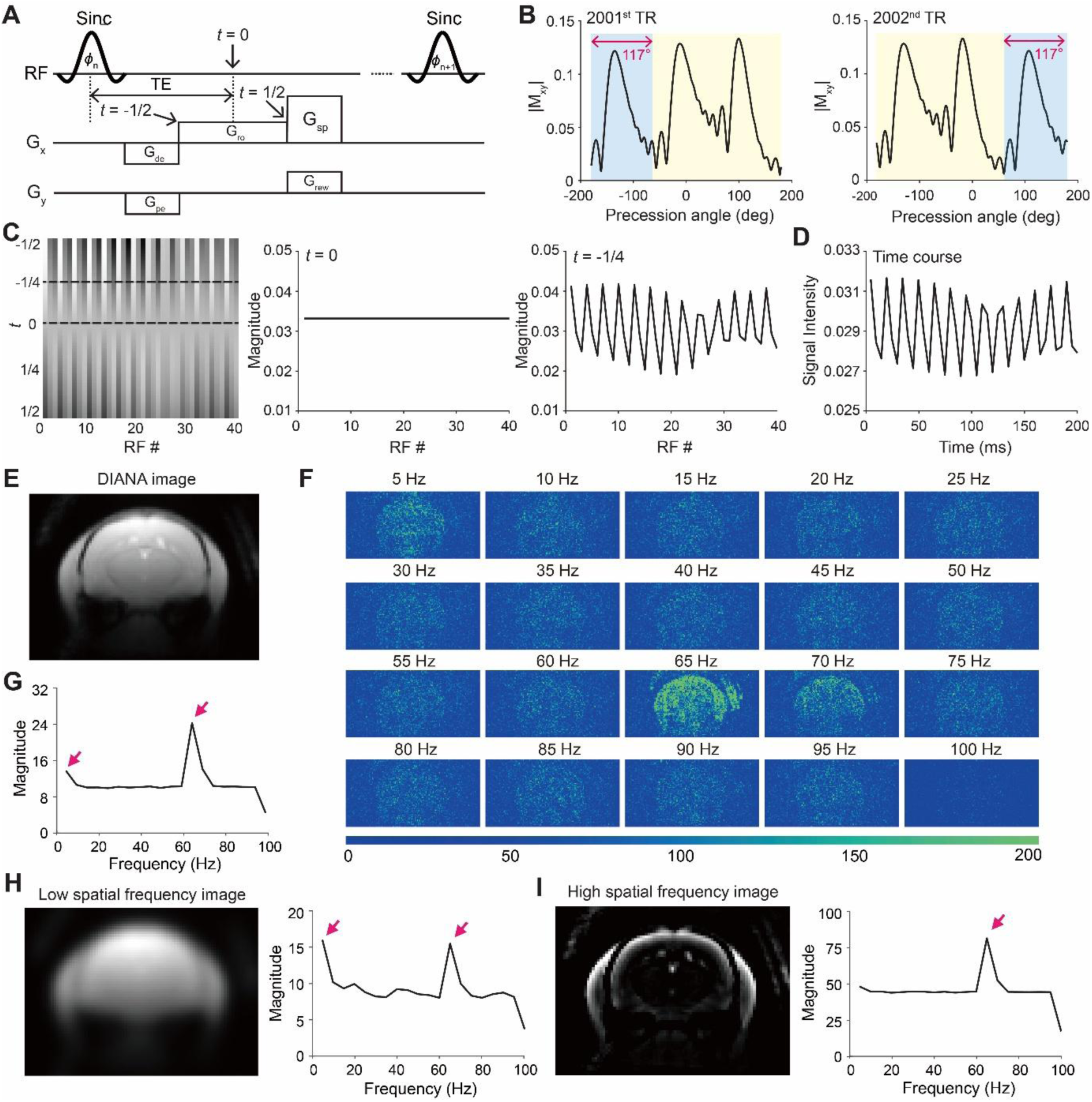
PSS oscillations due to RF spoiling in gradient-echo sequence. **(A)** Schematics of the gradient-echo pulse sequence including both RF and gradient spoiling. *t* is the normalized time for a readout gradient with a range of [-1/2, 1/2]. **(B)** Simulated magnetization profiles within a single voxel immediately following the 2001^st^ and 2002^nd^ excitation RF pulse when using an RF spoiling phase increment of 117°. **(C)** Concatenation of 40 echoes acquired from the 2001^st^ to 2040^th^ TR to visualize signal oscillations (left). Steady-state transverse magnetization at TE (middle) and signal oscillations near the edges of k-space (right) were shown. **(D)** Time series after 1D Fourier transform for a single voxel. **(E to G)** Spectral power density maps (E) for the averaged DIANA images (F) (*n* = 5 mice) and mean frequency spectrum averaged over all voxels (G). **(H)** Low spatial frequency image reconstructed using k-space around the center (left, *n* = 5 mice) and mean frequency spectrum (right). **(I)** High spatial frequency image reconstructed using the outer part of k-space (left, *n* = 5 mice) and mean frequency spectrum (right).

where *θ* is the total phase accumulated during TR (Fig. 3B).

Assuming that the spoiler gradient with the 2π gradient spoiling moment is applied only in the readout direction, the signal equation of a spoiled gradient-echo sequence at the *n*^th^ phase-encoding step is expressed as

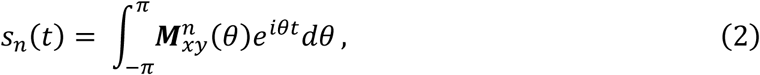

where *t* is the normalized time with the range of [-1/2, 1/2] during the readout gradient *G*_ro_ (see Fig. 3A). According to Eq. (2), *s*_n+1_(*t*) is not equal to *s*_n_(*t*) except when *t* = 0 (or TE). In other words, the signal is different at each phase-encoding step, but the signal is the same at TE because *θ* has a period of 2π and thus the integration results are the same. When RF spoiling is implemented at each phase-encoding step using a quadratic phase cycling scheme, there are two types of periodicity of **M***_xy_* in the phase-encoding direction, depending on whether the number of *ψ* rotations required for **M***_xy_* to return to its initial phase is an integer or not. If this number is an integer *n* for a given *ψ*, then **M***_xy_* repeats every *nψ.* For example, when *ψ* is set to 117°, **M***_xy_* becomes identical every 40 repetitions, as follows:

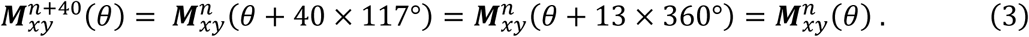

If the number of *ψ* rotations required for **M***_xy_* to return to its initial phase is a rational number, not an integer, **M***_xy_* becomes identical every 360°/117° (≍ 3.08) repetitions, as follows:

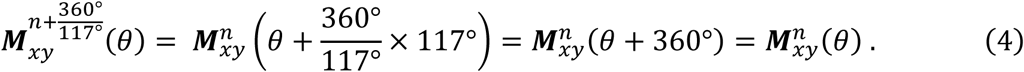

In this case, both periodic oscillations can cause ghosting artifacts along the phase-encoding direction in conventional gradient-echo images, which are shifted by *N*_y_/(360°/*ψ*), where *N*_y_ is the number of voxels along the phase-encoding direction.

To gain further insight into the behavior of spoiled gradient-echo signals with respect to RF spoiling, we also performed Bloch simulations. Gradient spoiling was initially implemented assuming a perfect gradient spoiling moment (= 2π). As shown in Fig. 3C, signal oscillation was observed along the phase-encoding direction at *t* = −1/4, whereas the signal was constant at *t* = 0. Moreover, ghosting artifacts shifted by 13 voxels (= 40 / (360°/117°) voxels) were observed along the phase-encoding direction (fig. S3A). However, in most practical applications, it is virtually impossible to achieve a perfect gradient spoiling moment of 2π due to hardware imperfections and eddy currents. Therefore, to get closer to the real situation, we slightly adjusted the gradient spoiling moment to 2.001π, and in this case, we observed weak oscillations even at *t* = 0 (fig. S3B).

### PSS oscillations due to RF spoiling in gradient-echo line-scan sequences

Considering the signal characteristics of gradient-echo sequences with RF spoiling discussed above, we investigated how PSS oscillations manifest themselves in gradient-echo line-scan sequences used in DIANA fMRI.

First, we investigated whether the PSS **M***_xy_* due to RF spoiling causes ghosting artifacts along the phase-encoding direction as described above in conventional gradient-echo imaging. According to Eq. (3), when the phase increment of RF spoiling is set to 117° (as in our mouse DIANA fMRI), the signals of all phase-encoding lines in each time frame become identical every 40 lines. Fortunately, when we performed the gradient-echo line-scan sequence in DIANA fMRI, we sampled 40 time points in each phase-encoding line, so ghosting artifacts due to PSS **M***_xy_* had little effect on the DIANA signal.

Next, we investigated whether the PSS **M***_xy_* due to RF spoiling has any effect on the time course of a gradient-echo line-scan sequence. According to Eqs. (3) and (4) in the previous section, since each phase increment of RF spoiling occurs in each phase encoding line and its periodicity exists in the phase encoding direction, PSS oscillations occur along the phase-encoding direction in conventional gradient-echo sequences. In contrast, PSS oscillations can occur along the time dimension in the gradient-echo line-scan sequence because each phase increment of RF spoiling occurs in each time frame of the time series (or each k-space line during the same phase encoding step) and thus its periodicity manifests in the time dimension (Fig. 3D). To confirm this effect *in vivo*, we reconstructed spectral maps for each frequency component via Fourier transform using group-averaged time-series images from DIANA fMRI at 11.7 T (*n* = 5 mice, Fig. 3E) and observed relatively strong 5 Hz signals within the mouse brain, and particularly strong 65 Hz and 70 Hz signals in the periphery of the brain (Fig. 3F and G).

Theoretically, these observations can be explained as follows: In a line-scan FLASH sequence using *ψ* of 117° in RF spoiling and TR of 5 ms, PSS **M***_xy_* can generate time series with frequency components of 5 Hz and 65 Hz, and sometimes even 70 Hz (6 Hz, 28 Hz, and 56 Hz in case of using *ψ* of 50° and the same TR). To elaborate, as shown in Eq. (3), the PSS **M***_xy_* is equalized every 40 TRs, so it can contribute to 5 Hz frequency component (i.e., 1 / (40 × TR) = 1 / 200 ms = 5 Hz) of the time series when using the gradient-echo sequence in a line-scan acquisition scheme. Similarly, according to Eq. (4), the PSS **M***_xy_* completes one rotation cycle every 360°/117°, which can contribute to a 65 Hz frequency component (i.e., 1 / (360°/117° × TR) ≍ 1 / 15.38 ms = 65 Hz) of the time series of the gradient-echo line-scan sequence, as shown in the spectral maps in Fig. 3F. A strong 70 Hz frequency component is also observed in Fig. 3F due to the harmonic frequency of 65 Hz, because according to the Nyquist theorem, 130 Hz (= 65 Hz × 2 = 130 Hz) is aliased to 70 Hz. Interestingly, Bloch simulations results further revealed that the 5 Hz signal oscillation is as strong as the 65 Hz signal oscillation around the center of k-space (fig. S4A), which represents the low spatial frequency components of the image, and the 65 Hz signal oscillation is strong in the outer part of k-space (fig. S4B), which represents the high spatial frequency components of the image.

Therefore, due to the combined effect of the low-frequency signal oscillations near the center of k-space observed in the Bloch simulation and the high tSNR obtained by placing the surface coil in the upper part of the mouse brain, it is reasonable to say that the 5 Hz frequency is relatively strong inside the brain, especially in the upper part of the brain, and that the 65 Hz frequency is strong around the periphery of the brain due to the high-frequency signal oscillations in the outer part of k-space as also observed in the Bloch simulation. Additionally, to verify that the simulation results are consistent with the *in vivo* data, we reconstructed images consisting of only low and high spatial frequencies, by applying a 2D Gaussian filter to the center of k-space with a full width at half maximum (FWHM) of 20 and 10 data points along the k_x_ and k_y_ directions, respectively. As a result, a 5 Hz component comparable to the 65 Hz component was observed in the low spatial frequency image (Fig. 3H), whereas only a strong 65 Hz component was observed in the high spatial frequency image (Fig. 3I).

### Frequency-offset dependence of PSS oscillations due to RF spoiling

Our next question was whether DIANA responses observed in VPL (∼23 ms) and S1FL (∼48 ms) at 7 T without trigger delay originated from neuronal activity rather than the PSS oscillations in a time series described above. To answer the question properly, it is particularly important to investigate how signal oscillations vary depending on several factors such as frequency-offset, spin-lattice relaxation time (T_1_), and T_2_.

First, we examined the frequency-offset dependence of the signal oscillations in the time series of a single voxel (Fig. 4A). If there is a static frequency-offset Δ*f*, Eq. (2) is expressed as

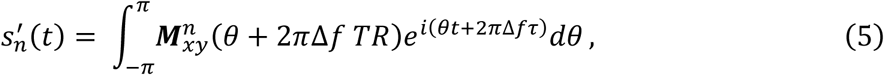

**Fig. 4.**
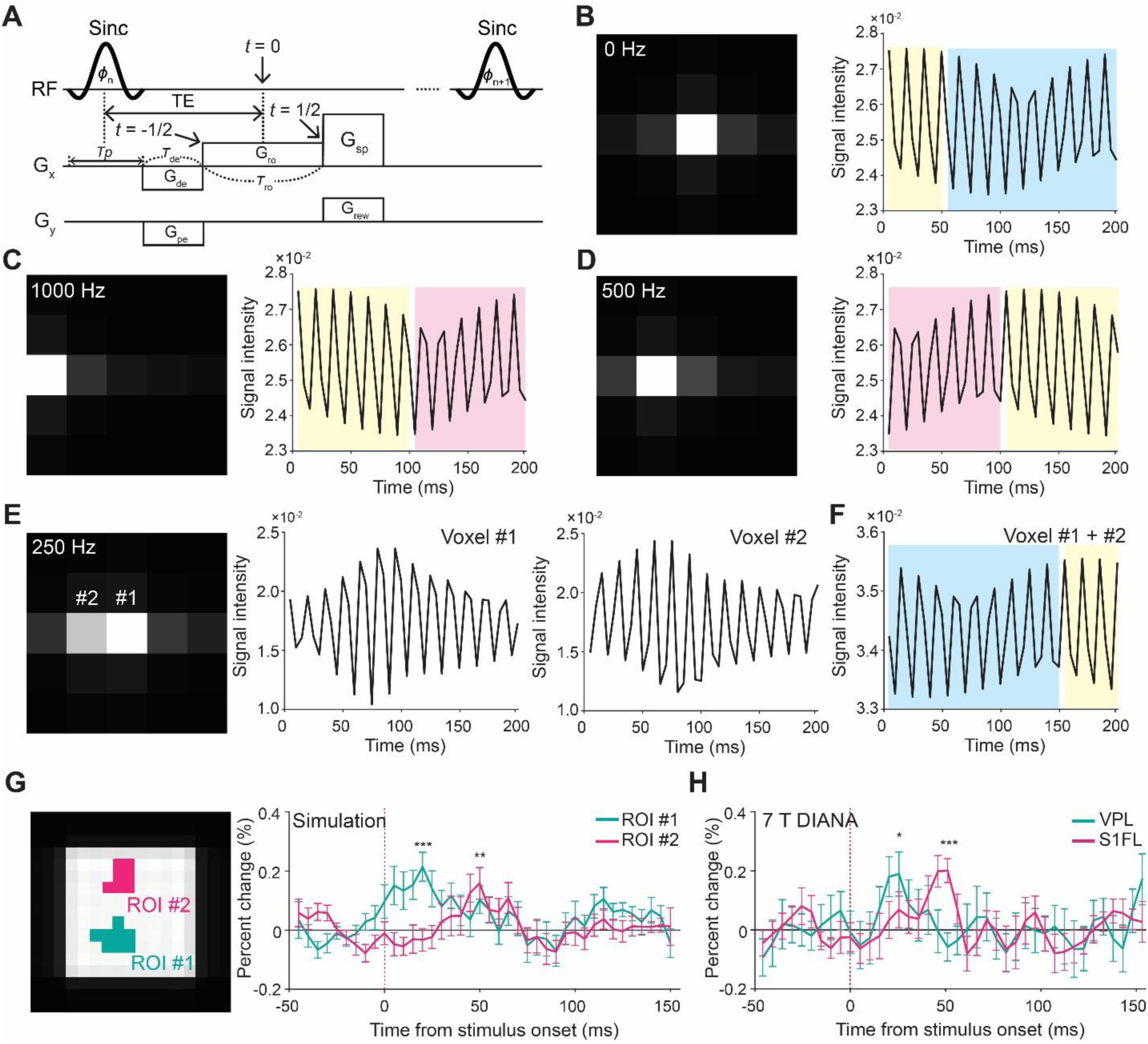
Effect of frequency-offset on PSS oscillations. **(A)** Schematics of the gradient-echo pulse sequence including both RF and gradient spoiling. *t* is the normalized time for a readout gradient with a range of [-1/2, 1/2]. **(B)** Reconstructed single voxel image and the time course without frequency-offset. **(C to D)** Reconstructed single voxel image (left) and the time course (right) with frequency-offset of 1000 Hz (C) and 500 Hz (D). Red and yellow shading indicates a shift in the time course by 20 time points. **(E)** Reconstructed single voxel image shifted by 0.5 voxel with a frequency-offset of 250 Hz (left) and the time course for each voxel (right, voxel #1 and #2). **(F)** The time course for the sum of the two voxels (voxel #1 + #2). Blue shading indicates a shift in the time course by 10 time points. **(G)** Selected ROIs (left) and the simulated time courses (right) extracted from the ROIs (*n* = 12). **(H)** DIANA responses obtained from the VPL (cyan) and S1FL (magenta) at 7 T (*n* = 12 mice). Vertical red dotted lines indicate the onset time of electrical forelimb stimulation. All data are mean ± SEM. *: *p* < 0.05, **: *p* < 0.01, ***: *p* < 0.001 for one-tailed Welch’s *t*-test.

where *τ* is the time from spin excitation. If Δ*f* TR is an integer, Eq. (5) can be simplified as

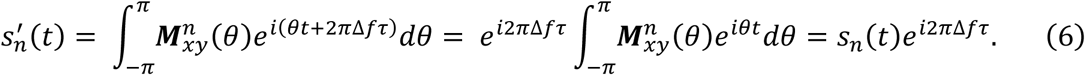

Substituting *τ = T_p_*/2 *+ τ*_de_ + *k*_x_ *τ*_ro_ Δ*x* (see Fig. 4A), where *T_p,_ τ*_de_ and *τ*_ro_ are the durations of RF pulse, pre-dephasing and rephasing readout gradient, respectively, and Δ*x* is the voxel size in the readout direction, Eq. (6) becomes

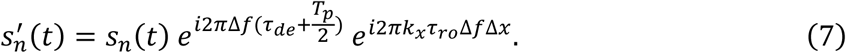

According to the Fourier shift theorem, Eq. (7) shows that the image is shifted by *τ*_ro_ Δ*f* Δ*x* voxels along the readout direction, that is,

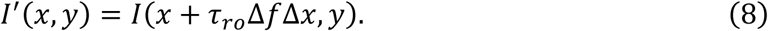

If Δ*f* TR is not an integer, Eq. (5) can be rewritten as follows by replacing *θ* + 2*π*Δ*f* TR with *θ*^′^:

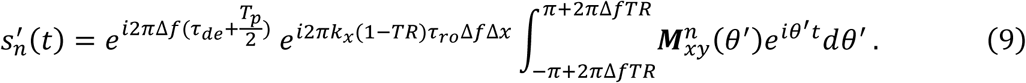

In Eq. (9), the second exponential term represents the shift in the image domain as in Eq. (8), and the last integral term represents the shift in the time domain.

As shown in Figs. 4B and C, assuming a 1000 Hz frequency-offset (Δ*f* TR = 5), the time course of a single voxel is maintained while its position is shifted by 2 voxels in the readout direction according to Eq. (8) (*τ*_ro_ Δ*f* = 2 ms × 1000 Hz = 2).

For a 500 Hz frequency-offset (Δ*f* TR = 2.5), the lower and upper bounds of the integration in Eq. (9) become [4π, 6π], which is in fact equal to [0, 2π] and can be simplified as

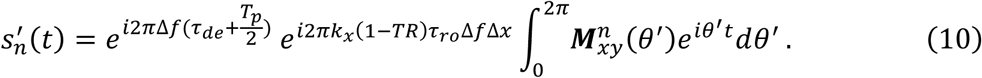

To examine how many time points are shifted from the original time course, recalling Eq. (2), the magnitude of the signal in (*n* + *k*)^th^ TR can be generalized as follows:

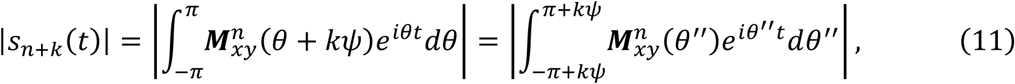

According to Eq. (11), a 117° RF spoiling phase increment (*ψ*) applied over 20 TRs (*k*) produces a total phase shift of π (*kψ* = 20 × 117° ≡ 180° mod 360°). Therefore, the time course described by Eq. (10) is equivalent to that of Eq. (2) shifted by 20 time points. Additionally, its position is shifted by 1 voxel in the readout direction in the same manner as Eq. (8) (*τ*_ro_ Δ*f* (1-TR) ≈ 2 ms × 500 Hz = 1, Fig. 4D).

For a 250 Hz frequency-offset (Δ*f* TR = 1.25), the single voxel signal is split into two voxels because its position is shifted by 0.5 voxel in the readout direction (*τ*_ro_ Δ*f* (1-TR) ≈ 2 ms × 250 Hz = 0.5, Fig. 4E). By adding the two voxel signals, we observed that the time course of the summed signal was shifted by 10 time points (10 × 117° ≡ 90° mod 360°, Fig. 4F).

So far, we have analyzed the frequency-offset dependence of PSS oscillations in the time series of a single voxel when using a gradient-echo sequence in line-scan format and have shown that the frequency-offset can lead to a shift in the time course in both the image domain and the time domain. These results suggest that in the mouse brain images acquired using a gradient-echo line-scan sequence, the time course of each voxel can represent a complex combination of temporally and spatially shifted signals from adjacent voxels when there are frequency-offsets, and that this temporal and spatial combination of signals may induce peak signals in the time courses at specific timing in specific regions.

To confirm this speculation, we further performed 2D Bloch simulations with 11×11 voxels (400 spins/voxel), assigning a frequency-offset to each spin from a Gaussian distribution with a mean of 0 Hz and a standard deviation of 20 Hz. To observe only the characteristics of the gradient-echo line-scan signal, no additional noise was introduced. As a result, we observed statistically significant peak responses similar to the DIANA responses observed in the VPL and S1FL at 7 T (∼0.19 %, 22.5 ± 1.91 ms, *: *p* < 0.05 for VPL and ∼0.20 %, 47.9 ± 1.99 ms, ***: *p* < 0.001 for S1FL, Welch’s *t*-test), with the similar peak amplitude and timing (∼0.21 %, 21.3 ± 2.62 ms, ***: *p* < 0.001 for ROI #1 and ∼0.16 %, 51.7 ± 2.91 ms, **: *p* < 0.01 for ROI #2, Welch’s *t*-test) (*n* = 12, Fig. 4G and H). These simulation results show that gradient-echo line-scan imaging with RF spoiling and without trigger delay can generate peak signals due to PSS oscillations at specific timing in specific regions, depending on frequency-offsets.

Additionally, we investigated the T_1_ and T_2_ dependence of the signal oscillations in a single voxel using Bloch simulation (fig. S5). T_1_ is a factor that affects only the magnitude of the steady-state signal in a gradient-echo line-scan sequence, which increases as T_1_ decreases. As expected, T_1_ affected only the overall magnitude of the signal oscillations and did not affect the frequency components of PSS oscillations (fig. S5A). In contrast to T_1_ dependence, T_2_ changes within relatively low T_2_ values had little effect on the overall magnitude and frequency components of PSS oscillations (fig. S5B). However, at high T_2_ values, new frequency components were introduced, which may be due to insufficient spoiling caused by the relatively high T_2_ compared to TR (fig. S5C).

### Various in vivo peak signals observed in functionally meaningful regions

According to the discussion so far, when RF spoiling is used, the gradient-echo line-scan sequence can generate spatially and temporally dependent peak signals due to imperfect steady state of **M***_xy_*, which is highly dependent on the frequency-offset. Also, according to *in vivo* mouse DIANA fMRI studies performed at different magnetic field strengths (e.g., 7 T and 11.7 T), DIANA signals appear to arise in the VPL and S1FL, with response times consistent with previous electrophysiological studies. In this section, considering the relatively wide spatial and temporal dependence of peak signals at various frequency-offsets, we further investigated whether these peak signals also appear in other regions of the mouse brain using data acquired using a FLASH line-scan sequence with RF spoiling (*ψ* = 117°) and trigger delay (25 μs) at 11.**7** T.

Using the spatiotemporal activation map (Fig. 1E) and the Allen Mouse Brain Atlas (*36*) to anatomically identify activation areas, we observed that peak signals with various timing occurred in several regions of the somatosensory network, including the posterior complex of the thalamic nucleus (PO), secondary somatosensory cortex (S2), secondary motor cortex (M2), and caudoputamen (CP), as well as in pain-related brain regions such as the mediodorsal nucleus of the thalamus (MD) and anterior cingulate cortex (ACC). The peak timing in some of these regions (e.g., dorsolateral VPL, S1FL and the middle part of S2) were found to be within a range similar to the timing of peak signals induced by trigger delay (∼30 ms), observed when averaging across all voxels in the brain, whereas the peak timing in other regions (e.g., M2 and dorsolateral POm) were outside this range (Fig. 5). All peak signal changes with various peak timing in these activation regions were statistically significant relative to the mean of the pre-stimulation period (fig. S6).

**Fig. 5.**
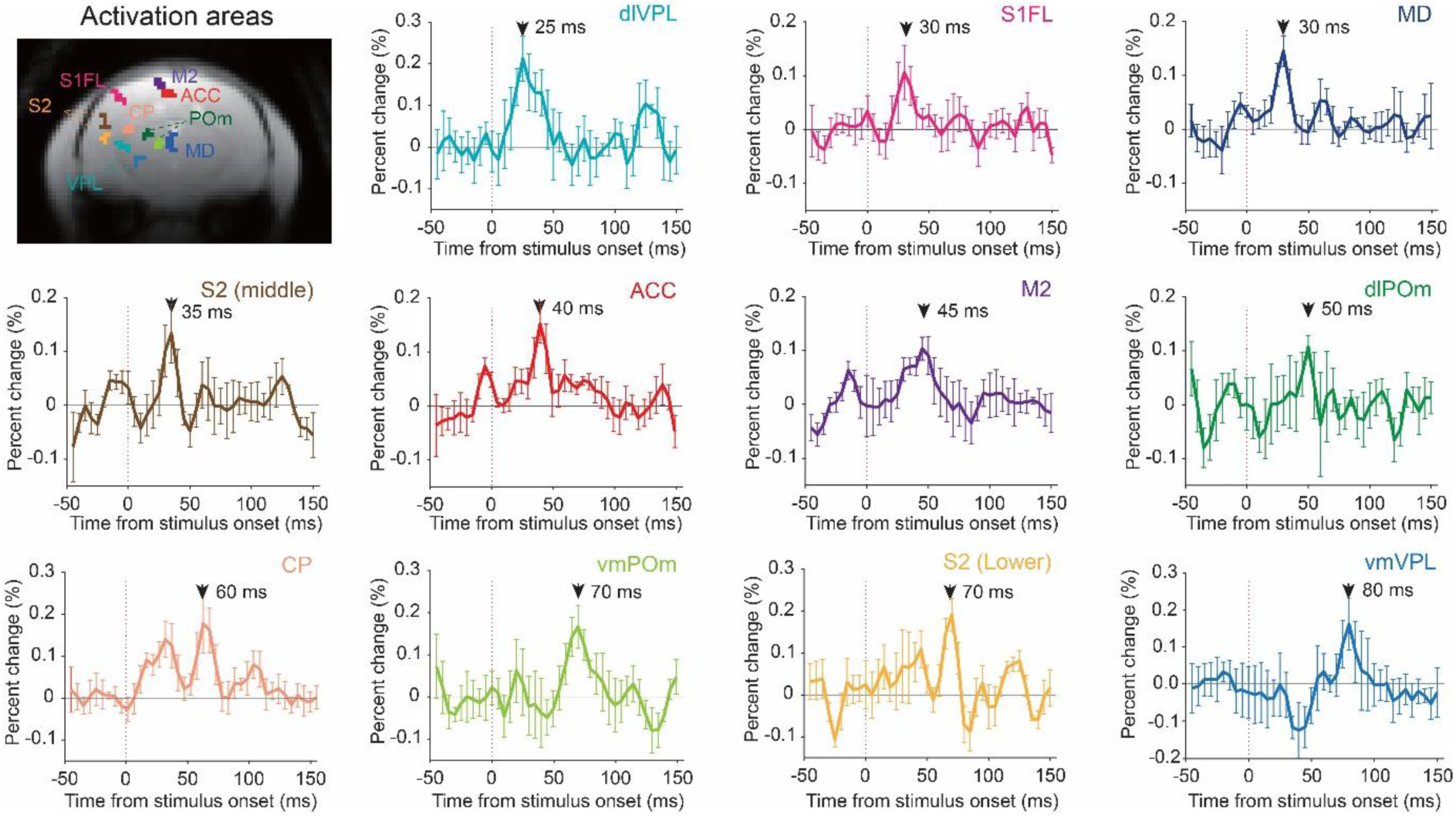
Various PSS-induced *in vivo* peak signals at 11.7. **T.** Activation areas associated with somatosensory network and pain-related circuits and DIANA time series with various peak timing (black arrowheads) in all activation areas are shown (*n* = 5 mice). Vertical red dotted lines indicate the onset time of electrical forelimb stimulation. All data are mean ± SEM.

It is noteworthy that the results identified several functionally known areas of the somatosensory network, in addition to VPL and S1FL areas that were mentioned in the previous sections. Nevertheless, it remains premature to interpret these peak signals as neuronal responses to sensory stimulation. This is because, at this stage, it is difficult to provide direct evidence to assert that the frequency-offset, which determines the timing and spatial positions of the peak signals and may arise from various physiological factors *in vivo*, is solely or dominantly attributed to neuronal activity. Further studies are required to link the frequency-offset (and T_1_, T_2_) dependence of **M***_xy_* under imperfect steady-state conditions to physiological factors that induce frequency-offset, including neuronal activity or hemodynamic effects during brain activation. When we performed 2D Bloch simulations repeatedly without trigger delay as in Fig. 4G, we could also observe statistically significant peak signals with a wide range of peak timing across various regions (fig. S7), due to the spatial and temporal dependence of PSS oscillations on frequency-offsets, because a frequency-offset (i.e., a Gaussian distribution with a mean of 0 Hz and a standard deviation of 20 Hz) was randomly assigned to each spin in each repetition.

## Discussion

In this study, we reported an update on the reproduction and interpretation of the DIANA signal. First, we reported the results of a replication experiment of the DIANA fMRI using forelimb electrical stimulation in medetomidine-anesthetized mice (*n* = 5 mice) at 11.7 T, including a novel data analysis method for creating spatiotemporal activation maps. This first replication experiment was performed to determine whether the results could be replicated using different stimulation areas (forelimb vs. whisker pad), different types of anesthetics (medetomidine vs. ketamine/xylazine), and different magnetic field strengths (11.7 T vs. 9.4 T) compared to the original study (*1*), but the experimental and sequence parameters were identical to the original experiment, including the application of RF spoiling (e.g., phase increment, 117°) and trigger delay (e.g., 25 μs). We observed sequential DIANA responses in the VPL and S1FL, with peak timing consistent with previous electrophysiological studies (*28–35*) and similar to those in the original study (*1*). In addition, we identified several functionally known areas of the somatosensory network at different peak timing through our proposed spatiotemporal activation mapping.

Next, since Phi Van et al. (*19*) reported unexpected peak signals due to trigger delay in their DIANA fMRI experiment and the timing of the trigger delay-induced peak signals was in a similar range to our original study (*1*), we investigated whether the DIANA response could persist even after removing the trigger delay. To this end, we implemented a trigger delay-removed sequence and performed DIANA fMRI without trigger delay using forelimb electrical stimulation in ketamine/xylazine-anesthetized mice (*n* = 12 mice) at 7 T. As a result, we also observed DIANA responses in the VPL and S1FL, with their peak timing consistent with previous electrophysiological studies (*28–35*). This suggests that DIANA fMRI still appears to be reproducible even in the absence of trigger delay effects.

However, in a recent human DIANA fMRI study, we discovered another important factor that significantly influences the DIANA signal and largely determines the spatial and temporal dependence of the signal, even in the absence of trigger delay. More specifically, during our analysis of human DIANA fMRI data, we found non-negligible frequency components (e.g., 6 Hz and 28 Hz) in the time series and recently discovered that these frequency components originate from the phase increase of RF spoiling in the gradient-echo line-scan sequence. This is because the phase increase of RF spoiling leads to a pseudo-steady state (abbreviated as PSS above) of **M***_xy_*, causing periodic oscillations of certain frequency components.

According to our investigation on the behavior of PSS oscillations based on theoretical analysis and Bloch simulations, the spatial location and temporal phase of PSS oscillations primarily depend on the frequency-offset. The temporal phase-shift caused by the frequency-offset in a single voxel may also help to intuitively understand the various timing of DIANA signals observed in various regions across the brain. However, in real-world situations, there may be limitations in accurately describing the specific locations and timing of peak signals due to PSS oscillations because the signal of a single voxel can be a combination of temporally and spatially shifted signals from adjacent voxels.

Several studies have reported periodic oscillations of specific frequencies in DIANA fMRI in both mice and humans. For example, Meng et al. (*10*) observed periodic fluctuations of ∼4 Hz in the mouse cortex when the interstimulus interval increased to 1,000 TR in DIANA fMRI, which they attributed to spontaneous firing of neurons. This interpretation seems reasonable because slow-wave activity in the range of 0.5 ∼ 4 Hz is dominant in anesthetized mice, and this is believed to originate from the neocortical neuronal network (*37*). However, our study suggests that these low-frequency oscillatory signals they observed are more likely to originate from PSS oscillations due to RF spoiling rather than spontaneous firing of neurons. As shown in Fig. 3, using RF spoiling with a phase increment of 117°, low-frequency oscillations of 5 Hz can be induced by PSS oscillations, particularly in the low spatial-frequency region. In some human DIANA fMRI studies, oscillatory signals with frequency components of 6 Hz, 28 Hz and 56 Hz were also observed (*7*, *38*), which could also be well predicted by PSS oscillations when the phase increment of RF spoiling was set to 50°, as in the Siemens human scanner. In summary, our recent study demonstrated that PSS oscillations due to RF spoiling, which are spatially and temporally dependent on frequency-offset, can significantly contribute to the generation of the DIANA signal.

Together with the previous findings of Phi Van et al. (*19*), it is concluded that DIANA signals can be significantly influenced by two important factors of the gradient-echo sequence: trigger delay and RF spoiling. Fundamentally, the physical mechanisms by which these two factors contribute to the DIANA signal can be explained in terms of imperfect steady state of magnetization. In other words, the trigger delay disrupts the steady state of magnetization of the spin system, while the phase increase of RF spoiling leads to a PSS of magnetization, which in turn causes PSS oscillations.

If RF spoiling is not used, the signal evolves toward a steady state where the magnetization (e.g., **M***_z_* and **M***_xy_*) converge to a constant value as the TR is repeated. In this case, the peak signals induced by the trigger delay occur because the magnetization undergoes additional recovery during the trigger delay, and the peak signals appear in the TR following the TR in which the trigger is applied. These peak signals can be significantly suppressed, when RF spoiling is applied, T_2_ is relatively short, and the magnetic field is relatively uniform (*19*).

With RF spoiling, the magnetization reaches a PSS rather than a perfect steady state, resulting in a periodic oscillatory behavior (*20*). The resulting PSS oscillations have spatial and temporal dependence on frequency-offset, which affects the spatial location and temporal shift of the peak signals in DIANA fMRI. Unlike the peak signals due to trigger delay, the peak signals due to PSS oscillations are not suppressed by changes in T_2_ and B_0_ inhomogeneity.

If both trigger delay and RF spoiling are present together, the trigger delay disrupts the PSS (not perfect steady state) of the magnetization and slightly modulates the behavior of the PSS oscillations, resulting in a signal with a mixture of the effects of these two factors. Interestingly, as described in Eq. (2), PSS oscillations are prominent in the outer part of k-space and diminish near the center of k-space (and, in principle, completely vanish at TE), such that the peak signals due to PSS oscillations dominate the high spatial-frequency regions. In contrast, since PSS oscillations are weakened near the center of k-space, the trigger-delay effect appears to be more pronounced near the center of k-space and thus dominates in the low spatial-frequency regions. To further validate these statements, we generated peak response time maps via spatiotemporal activation mapping, using low and high spatial-frequency images reconstructed from 11.7 T mouse data. As expected, in the low spatial-frequency image (*n* = 5 mice), peak signals with nearly identical timing were observed across the brain, likely reflecting the trigger delay (fig. S8A), whereas in the high spatial-frequency image (*n* = 5 mice), the timing of peak signals varied substantially, likely due to PSS oscillations influenced by B_0_ inhomogeneity (fig. S8B). These results also suggest that PSS oscillations tend to be more pronounced in smaller ROIs, but as ROI size increases, their relative contribution decreases, making the trigger-delay effect more pronounced.

Finally, the frequency-offset on which PSS oscillations primarily depend, both spatially and temporally, is conceptually equivalent to the variation in the main magnetic field (or B_0_ inhomogeneity) in MRI. *In vivo*, the frequency-offset is determined not only by the time-invariant differences in tissue susceptibility, but also by any physiological variations that can induce B_0_ inhomogeneity, including neuronal activity and hemodynamic effects such as those underlying the BOLD response. Therefore, if PSS oscillations are the main cause of DIANA signals, it may be premature to interpret DIANA signals as neuronal responses to sensory stimulation until further research can explain how the frequency-offset dependence of PSS oscillations is primarily dependent on neuronal activity in DIANA fMRI. In this context, follow-up research worth pursuing would be to investigate what can be observed in DIANA fMRI without PSS oscillations. One possible way to effectively suppress PSS oscillations may be to refer to a previously proposed method used to suppress ghost artifacts in FLASH images (*22*), so that it can be applied to a line-scan imaging strategy.

In some respects, peak signals arising from PSS oscillations may be regarded as artifacts that contaminate DIANA signals and need to be eliminated. However, this perspective does not readily explain our observation that the DIANA fMRI results discussed here identified several functionally known areas of the somatosensory network, including the VPL and S1FL, and the timing of the DIANA responses was consistent with previous electrophysiological studies, particularly in VPL and S1FL (e.g. at 7 T, 9.4T, and 11.7 T). The observed consistency of DIANA fMRI results across activated regions and response timing at different magnetic field strengths is difficult to ascribe to coincidence alone. Therefore, we anticipate that sufficient time and comprehensive support from the academic community would be required to fully elucidate the relationship between DIANA signals and brain activation, including neuronal activity.

## Materials and Methods

### 7 T MRI experiments

MRI experiments were conducted using a 72/112 mm inner/outer-diameter volume coil for RF transmission and 2×2 surface array coil for signal reception on a 7 T animal scanner (BioSpec 70/20 USR, Bruker, Ettlingen, Germany).

#### Animals

Twelve iDTR transgenic mice (male, 23 – 35 g, 5 – 12 months old, originate from Korea Research Institute of Biosience and Technology, South Korea) were used in the experiments. All procedures were approved by the Institutional Animal Care and Use Committee of Sungkyunkwan University (SKKUIACUC-2023-11-02-1) and National Cancer Center (NCC-22-855-006). Mice were housed under standard conditions, and food and water were provided *ad libitum*.

#### Animal preparation

Mouse were initially anesthetized with 4% isoflurane for 2 minutes in an induction chamber. Subsequently, a mixture of ketamine (100 mg/kg) and xylazine (10 mg/kg) was applied intraperitoneally. After an hour, additional doses with mixture of ketamine (25 mg/kg) and xylazine (1.25 mg/kg) were applied every 40 minutes or whenever the respiration rate exceeded 180 bpm. An injection tube connected to a syringe positioned outside the magnet bore was used to deliver additional dose during the experiment without disturbing the setup. Animals were fixed on an animal cradle using ear bars and a tooth bar. Rectal temperature was maintained between 35–37°C using an air heating system. Respiration rate (target range: 120–180 breaths per minute) was monitored using a pneumatic pressure sensor (Model 1030, Small Animal Instruments, USA). For electrical stimulation, two needle electrodes were subcutaneously inserted into the interdigital spaces between the second and third digits, and between the third and fourth digits of the right forepaw.

#### Reference imaging

To visualize anatomical brain structures and check the location of the primary somatosensory cortex (S1) and the sensory thalamic nuclei, ten 0.5 mm thick coronal slices were acquired using a 2D rapid acquisition with refocusing echoes (RARE) sequence. The scan parameters were as follows: TR/TE = 2000/12 ms; FOV = 16 × 16 mm²; matrix size = 256 × 256; and effective TE = 36 ms. After identifying coronal slices containing S1 and sensory thalamic nuclei, we acquired two 1 mm thick coronal slices covering S1 and sensory thalamic nuclei, respectively. The anterior slice was positioned to cover S1, and the posterior slice was positioned to cover the sensory thalamic nuclei.

#### DIANA fMRI

Functional imaging was performed using a 2D FLASH-based line-scan imaging sequence (ParaVision 5) after removing trigger-related delay. Scan parameters were as follows: TR/TE = 5/2 ms; FA = 4.6°; FOV = 16 × 16 mm²; matrix size = 72 × 72; slice thickness = 1 mm; and scan time, 18 s/trial. Sufficient dummy scans (10s, 2000 TR) were used to achieve steady-state magnetization prior to the main acquisitions. Both gradient and RF spoiling were used to suppress the effects of residual transverse magnetization. RF spoiling was applied by default module in Bruker Paravision 5 software, that is, using a quadratic phase cycling schedule to increment the phase of *n*^th^ excitation RF pulse by *n* × 117°. Therefore, the phase of *n*^th^ excitation RF pulse *ϕ*_n_ was given by

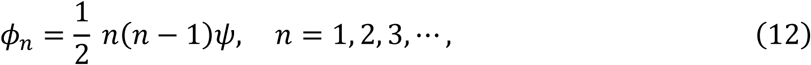

where *ψ* = 117°. In this case, the phase of the RF pulse increases at each k-space line of the time series within the interstimulus interval, whereas for conventional RF spoiling it increases at each phase encoding step. An identical two 1 mm thick slices acquired with 2D RARE sequence were acquired for DIANA fMRI. Electrical stimulation of 1.0 mA current strength, 1 ms pulse duration, and 4 Hz frequency was delivered to the right forepaw during the scan, using stainless steel electrodes connected to the isolated pulse stimulator (A-M System Model 2100, USA). The stimulus was repeated 72 times, corresponding to the number of phase-encoding steps with an interstimulus interval of 250 ms. The stimulation paradigm consisted of 100 ms pre-stimulation, 1 ms stimulation, and 149 ms post-stimulation. For each mouse, 40 trials were acquired. To minimize neural adaptation, a 90-second rest period was provided every five trials.

#### Sequence Modification

A 2D FLASH-based line-scan imaging sequence for DIANA fMRI was modified to remove trigger-related delay in Bruker’s ParaVision 5 pulse programming environment. Within the Bruker ‘ppg’ pulse sequence code, a loop counter variable ‘lTrig’ was initialized to zero. The trigger signal was sent externally via ‘TTL1’ channel using ‘TTL1_HIGH’ and ‘TTL1_LOW’ commands, which toggle the transistor-transistor logic (TTL) signal depending on the current value of ‘lTrig’. Each phase-encoding line consisted of segmented loops total of 50 TRs. The number of segments in each phase-encoding line was specified by ‘NTrigSegments’, and the number of TR within each segment was specified by ‘trigSegment’. Originally, NTrigSegments’ was set to 2 and ‘trigSegment’ was set to [1, 49], meaning that the first segment contained one TR loop, and the second segment contained 49 TR loops. The outer loop iterated over the number of segments defined by NTrigSegments, and the inner loop iterated according to the corresponding value of trigSegment. A delay was introduced when transitioning from the inner loop to the next iteration of the outer loop. Specifically, a trigger-related delay of 3 µs was introduced at the beginning of the first and second segment, which correspond to the beginning of the first and second TR of each phase-encoding line (1 µs from the start loop, 1 µs from the TTL toggle, and 1 µs from the increment operation). Therefore, total trigger-related delay of 6 µs was introduced for each phase-encoding step (3 µs/segment × 2 segments = 6 µs). To eliminate the trigger-related delay, we modified both trigger delays of 3µs to 0µs and moved the ‘TTL1_HIGH’ and ‘TTL1_LOW’ commands into the inner loop, ensuring that each TR maintains 5 ms. The loop structure was also adjusted by setting ‘NTrigSegments’ to 1 and ‘trigSegment’ to [50], so that all 50 inner loops were executed within a single outer loop. This approach ensures uniform timing across all TRs and eliminates the trigger-related delays previously introduced by loop-dependent operations.

#### Phantom imaging

Phantom imaging was performed using a 2D FLASH-based line-scan imaging sequences (ParaVision 5) with a 15 mL conical tube (diameter: 1.7 cm) filled with saline solution. We modified the second delay introduced at the beginning of the second segment, which correspond to the beginning of the second TR, to 52 µs, 14 µs, and 3 µs, respectively, and implemented the modified sequences in the MR scanner to verify the effect of the trigger-related delay. Scan parameters were as follows: TR/TE = 5/2 ms; FA = 4.6°; FOV = 16 × 16 mm²; matrix size = 72 × 72; and slice thickness = 1 mm. 50 trials were acquired for each modified sequence.

### 11.7 T MRI experiments

All experiments for DIANA and BOLD fMRI were performed using a surface cryoprobe on a 11.7 T animal scanner (BioSpec 117/16 USR, Bruker, Ettlingen, Germany).

#### Animals

Six C57BL/6J wild-type mice (29 ± 4 g, 3 – 3.5 month old, Charles River Laboratory, Saint-Germain-Nuelles, France) were used in the experiments. This study was carried out with approval by the Institutional Animal Care and Use Committee (APAFIS#18001-2018121010505627 v1).

#### Animal preparation

Mice were initially anesthetized with 4% isoflurane and maintained at 2% during animal preparation and were placed prone on a heat-regulated cradle. Xylocaine droplets were applied in the ears as a local anesthetic, and ear bars and a tooth bar were used to secure the animal’s head still in a nose cone. The mice were self-breathing with a continuous supply of oxygen and air gases (1:1 ratio) through the nose cone at a rate of 1 liter/min. A subcutaneous catheter was placed in the back to deliver a 0.3 mg/kg bolus of medetomidine (Domitor®, Vetoquinol) followed by a continuous infusion at 0.6 mg/kg/h. The isoflurane level was slowly reduced over 5 min until complete discontinuation. Rectal temperature (37°C) and respiration rate (140 – 160 bpm) were controlled with the SAII’s MR-compatible Model 1030 Small Animal Monitoring and Gating System (SA Instrument, Inc., Stony Brook, NY, USA).

#### BOLD fMRI

2D gradient-echo echo planar imaging (EPI) was used for BOLD fMRI. Scan parameters were as follows: TR/TE = 1000/15 ms; FA = 45°; FOV = 24 × 12 mm^2^; matrix size = 128 × 64; dummy scans = 40 TR; and slice thickness = 0.5 mm or 0.8 mm. First, to check the location of sensory thalamic areas and S1FL, ten 0.5 mm thick coronal slices including the thalamus and S1FL were acquired. Electrical stimulation of 0.5 mA current strength, 2 ms pulse duration, and 10 Hz frequency was delivered to the right forepaw during the scan, using stainless steel electrodes connected to the isolated pulse stimulator (A-M System Model 2100). The stimulation paradigm consisted of 5 blocks of 20s-on/20s-off cycles. When examining which coronal slices the thalamus and S1FL responded to, BOLD responses in the S1FL and thalamus were approximately 4 to 5 coronal slices apart. Based on these results, a single 0.8 mm thick oblique slice was set using the same stimulation paradigm to observe BOLD responses in the thalamus and S1FL simultaneously. The oblique slice was acquired before and after BOLD fMRI experiments to determine whether or not the mouse responded to the stimulation during the DIANA fMRI experiment. A total of 3 runs were repeated.

#### DIANA fMRI

Experiments were performed using 2D FLASH based line-scan imaging sequence (ParaVision 6.0.1). Scan parameters were as follows: TR/TE = 5/2 ms; FA = 4°; FOV = 25.6 × 12.8 mm^2^; matrix size = 128 × 64, slice thickness = 0.8 mm; bandwidth = 75 kHz; and scan time = 12.8 s/trial. Sufficient dummy scans (8 s, 1600 TR) were used to achieve steady-state magnetization prior to the main sequence. Both gradient and RF spoiling were used to suppress the effects of residual transverse magnetization. RF spoiling was applied by default module, which is identical to Eq. (12). An 0.8 mm thick oblique slice was acquired for DIANA fMRI with 0.5 mA current strength, 1 ms pulse duration, and 5 Hz frequency. The stimulus was repeated 64 times, corresponding to the number of phase-encoding steps with an interstimulus interval of 200 ms. The stimulation paradigm consisted of 50 ms pre-stimulation, 1 ms stimulation, and 149 ms post-stimulation. 40 trials per mouse were acquired and used for analysis. To minimize neural adaptation, a 120-second rest period was provided every five trials.

### fMRI data processing

All DIANA fMRI data were processed using home-built MATLAB (R2022b, MathWorks), Analysis of Functional Neuroimages package (AFNI) (*39*), and FMRIB Software Library (FSL) (*40*). All analyses of DIANA fMRI data were referenced to the Allen Mouse Brain Atlas (*36*) (Allen Brain Institute, http://mouse.brain-map.org).

#### BOLD fMRI

Individual BOLD activation maps were generated using preprocessing and a general linear model (GLM) analysis with 5 boxcar blocks of 20s-on/20s-off cycles after averaging 3 runs. Preprocessing steps were as follows: linear detrending to remove signal drift and time course normalization relative to the first image. After the preprocessing and GLM analysis, individual BOLD activation maps were generated with a statistical threshold of uncorrected p < 0.05 and cluster size > 5 and overlaid on the EPI images. Additional spatial smoothing was not used to generate BOLD activation maps. Time courses were extracted from circular ROIs of ∼0.6 mm (3.2 pixels) radius in the thalamus and S1FL regions.

#### DIANA fMRI

For group analysis, the trial-averaged DIANA image of each mouse was linearly registered to the trial-averaged DIANA image of mouse #1 using FMRIB’s Linear Image Registration Tool (FLIRT) function in FSL. Transformation matrix for each mouse was extracted by linearly registering the DIANA images averaged across all 40 frames and 40 trials for each mouse to the DIANA image of mouse #1 averaged across all 40 frames and 40 trials, using tri-linear interpolation. Next, each transformation matrix was applied to all 40 frames and 40 trials for corresponding mouse using ‘applyxfm’ function in FSL.

Spatiotemporal activation mapping was performed to obtain DIANA-activated specific brain regions. Prior to spatiotemporal activation mapping, several preprocessing steps were taken: Voxel-wise temporal smoothing was first applied using a three-point Gaussian kernel, and each frame was spatially smoothed with a Gaussian kernel of ∼0.5 mm FWHM instead of ROI averaging. After that, time series in each voxel was normalized relative to the mean of the pre-stimulation period, and linear detrending was applied to the time series to remove signal drift. Finally, voxels with at least one data point with z-score > 3 or < −3 were excluded as outliers, considering the histogram of all data points in the time series of all voxels. These outlier voxels were highly concentrated in the lower part of the brain, where the tSNR is relatively low because a surface coil was placed on top of the brain. After the preprocessing, a peak response time map was generated by calculating the time that has maximum amplitude of the time series in each voxel within the brain. Next, spatiotemporal clustering was performed by clustering the voxels with the same peak response time and selecting clusters with size greater than or equal to 4 for each time frame. Then, to test the statistical significance of the DIANA response in each cluster, statistical comparisons were made between the baseline mean of the pre-stimulus period and every post-stimulus time point in the average time series for each cluster using data from all five rats. Through this statistical process, clusters with at least one statistically significant data point survived to form the spatiotemporal activation map. Using the spatiotemporal activation map, activation voxels as clusters were defined in the forelimb sensory processing regions of interest. Finally, time series were extracted from those defined ROIs using temporal smoothing using a three-point Gaussian kernel, ROI averaging instead of spatial smoothing, signal normalization, and linear detrending. For 7 T DIANA fMRI, the 11^th^ to 20^th^ TRs were used as baseline, excluding the first 10 time points and ROI was simply defined by a 3 × 3 square. The DIANA response latency for each mouse was calculated for the maximum peak within a 45 ms window size, from 20 ms before to 25 ms after the time of the maximum peak of the average time series.

The spectral amplitude maps were generated for each frequency component by calculating the amplitude of frequency component for each voxel. The amplitude was computed by fast Fourier transform. The images consisting of only low and high spatial frequencies were reconstructed, by applying a 2D Gaussian filter centered at the center of k-space and with a FWHM of 20 and 10 data points along the k_x_ and k_y_ directions, respectively.

### Bloch simulation

1D and 2D Bloch simulations were performed to investigate the signal behavior of a FLASH-based line-scan imaging sequence. The simulations were performed using home-built MATLAB code (R2022b, MathWorks) and executed on a GPU-enabled system for computational efficiency. Scan parameters used in Bloch simulations were as follows: TR/TE = 5/2 ms; FA = 4°; and matrix size = 72×54 for the 2D Bloch simulation; and matrix size = 72×1 for the 1D Bloch simulation.

RF excitation was simulated by assuming that the isochromat experiences an instantaneous rotation of the flip angle at the center of RF pulse. RF spoiling was implemented using a quadratic phase cycling with a phase increment of 117°, following the standard formulation of Eq. (13).

The spins experienced spatially varying phase rotations due to the applied gradient along the frequency-encoding direction. A dephasing gradient is applied immediately after RF excitation, followed by the readout gradient during signal acquisition. The dephasing gradient was simulated by dephasing the spins of each voxel by -π and the time-resolved readout gradient was simulated across 72 time points to simulate the temporal evolution during each acquisition window. The spins experienced total 2π rephasing per voxel during the readout gradient. To spatially encode along the phase-encoding direction, a gradient is applied at each TR with a fixed amplitude corresponding to the desired k-space line. After signal acquisition, a phase-rewinder gradient is applied to undo the accumulated phase from the phase-encoding gradient. The rewinder has the same area but opposite polarity, effectively resetting the net phase accumulation in ideal conditions. Both the phase-encoding and rewinder gradients were implemented with a 2π dephasing and rephasing per voxel of the spins in the transverse plane, respectively. For 1D Bloch simulations, the phase-encoding and rewinder gradients were omitted. Finally, to eliminate residual transverse magnetization, a spoiler gradient was applied along the frequency-encoding direction at the end of each TR, introducing a π phase shift per voxel, leading to a total dephasing of 2π within the single voxel.

The simulation included 2,000 dummy scans to achieve steady-state magnetization, followed by 2,160 acquisitions (54 phase-encoding lines × 40 time points) for 2D and 40 acquisitions for 1D, mimicking the full line-scan imaging process. Tissue relaxation parameters were defined as T_1_ = 2000 ms and T_2_ = 35 ms for all spins. To incorporate B_0_ inhomogeneity, frequency-offset from the Gaussian distribution with a mean of 0 Hz and a standard deviation of 20 Hz was assigned for each spin, and their effects on phase accumulation during each gradient lobe (dephasing, readout, and spoiler) were simulated.

Each 2D Bloch simulation computed a single voxel containing 100 spins along the frequency-encoding and phase-encoding direction, respectively, for a total of 10,000 spins, or 11×11 voxels containing 20 spins along the frequency-encoding and phase-encoding direction for each voxel, respectively, for a total of 48,400 spins. For 1D Bloch simulation, 100 spins were computed. The transverse magnetization was sampled during readout, and complex-valued time series data were computed by summing the signals across all spins. The final k-space data were reshaped into a 3D matrix (time points × phase-encoding × readout) and 2D Fourier transformed along phase-encoding and readout directions to generate magnitude images. This full process was repeated for 40 trials per run.

The time series were extracted from the defined ROIs identified by spatiotemporal activation mapping, and temporal smoothing was performed using a three-point Gaussian kernel, ROI averaging instead of spatial smoothing, signal normalization, and linear detrending. The time series of single voxels were extracted without any processing.

### Statistics

For all statistical comparisons, Shapiro-Wilk test was performed to test normality except for samples not less than 30. The samples with more than 30 were considered to follow a normal distribution based on the central limit theorem (*41*). For all statistical comparisons between the two groups satisfying normality, a Levene’s test was performed to check for homoscedasticity, followed by a one-tailed paired t-test or Welch’s t-test. If normality was not satisfied in this case, a one-tailed Wilcoxon signed-rank test was performed. Quantitative data were all expressed as mean ± standard error of the mean (SEM) or mean ± standard deviation (SD).

## Acknowledgments

We thank J. Valette and C. Baligand (MIRCen, CEA) for arranging the experiments at 11.7 T and for valuable scientific discussion on data acquisition and analysis; S. Malaquin, C. Hery, and E. Mougel (MIRCen, CEA) for preparing and helping experiments. We also thank M. Lowe, W. Shin, A. Nemani, K. Sakaie (Imaging Institute, Cleveland Clinic) for insightful statistical and scientific discussions.

## Author contributions

Conceptualization: J.-Y.K. Methodology: J.-Y.K., S.P., and J.-Y.P.

Investigation: J.-Y.K., S.P., H.C., and D.K. Visualization: J.-Y.K. and S.P.

Project administration: J.-Y.P. Supervision: J.-Y.P.

Writing – original draft: J.-Y.K.

Writing – review & editing: J.-Y.K. and J.-Y.P.

## Data and materials availability

All data needed to evaluate the conclusions in the paper are present in the paper and/or the Supplementary Materials.

## Supplementary Materials

Figs. S1 to S8

