## Supplementary figures for "Update on the reproduction and interpretation of DIANA fMRI"

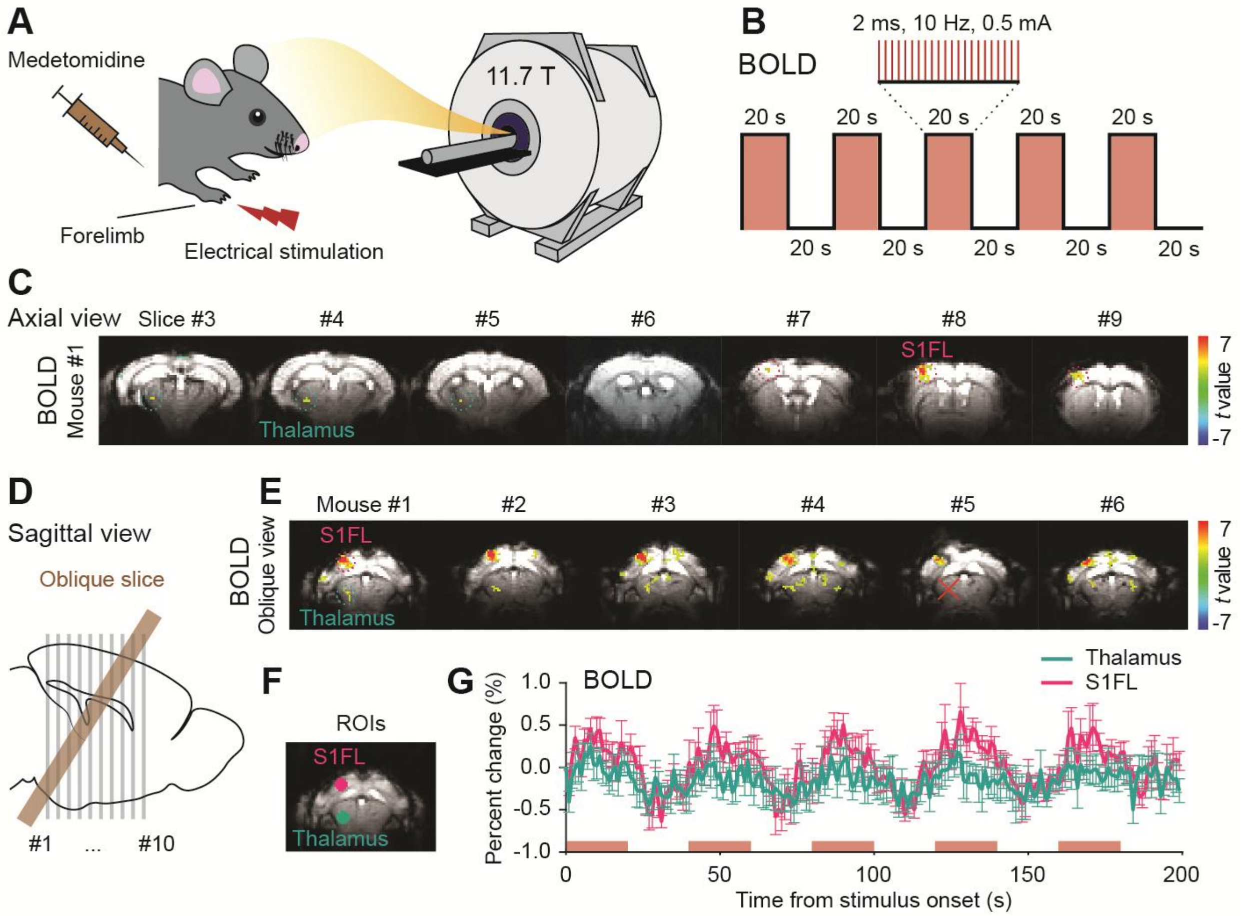


**Fig. S1.**

**BOLD fMRI in medetomidine-anesthetized mice at 11.7 T. (A to B)** Schematics of the BOLD fMRI experiment (A) and stimulation paradigm (B) to capture BOLD responses to electrical right forelimb stimulation. **(C)** BOLD responses in coronal slices of a representative 5 mouse (mouse #1). **(D)** Location of the set oblique slice in sagittal plane. **(E)** BOLD responses from 6 mice in the oblique slice. **(F)** Circular ROIs selected for the thalamus (cyan) and S1FL (magenta). **(G)** BOLD time courses extracted from the ROIs of the thalamus and S1FL ($n$ = 5 mice). Red box areas indicate the 20s-on state of electrical forelimb stimulation (G). All data are mean ± SEM.

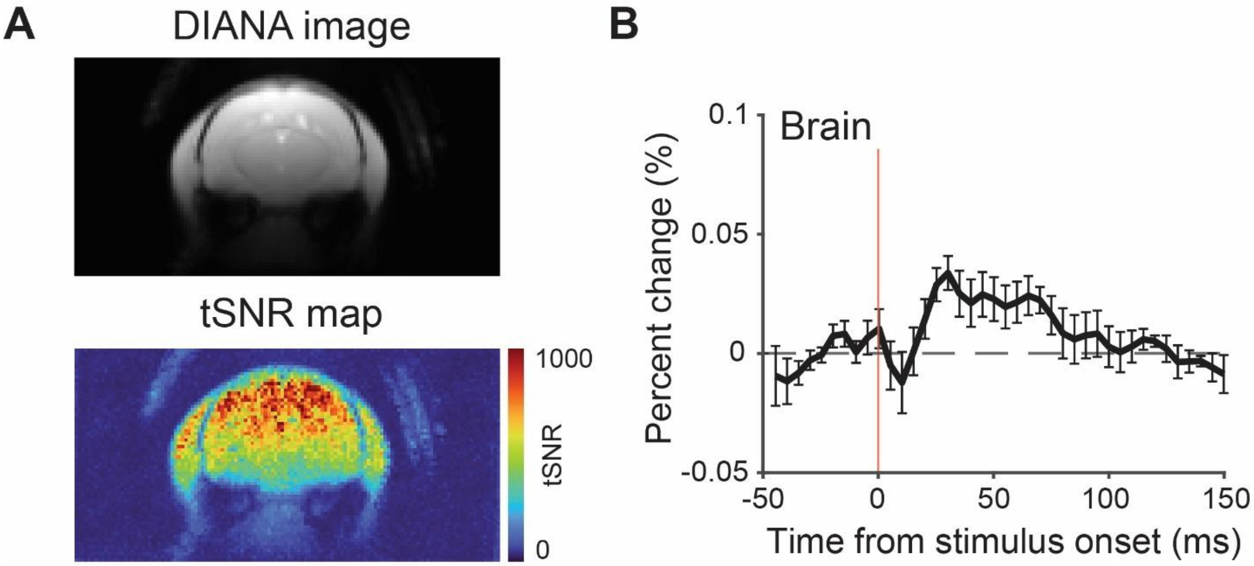


**Fig. S2.**

**Trigger delay *in vivo* peak signal in 11.7 T DIANA fMRI (A)** Averaged DIANA image for 200 trials (= 5 mice × 40 trials/mouse) (top) and tSNR map (bottom). **(B)** Trigger delay *in vivo* peak signal within the mouse brain (black, *n* = 5 mice). Vertical red lines indicate the virtual onset time of electrical forelimb stimulation. All data are mean ± SEM.

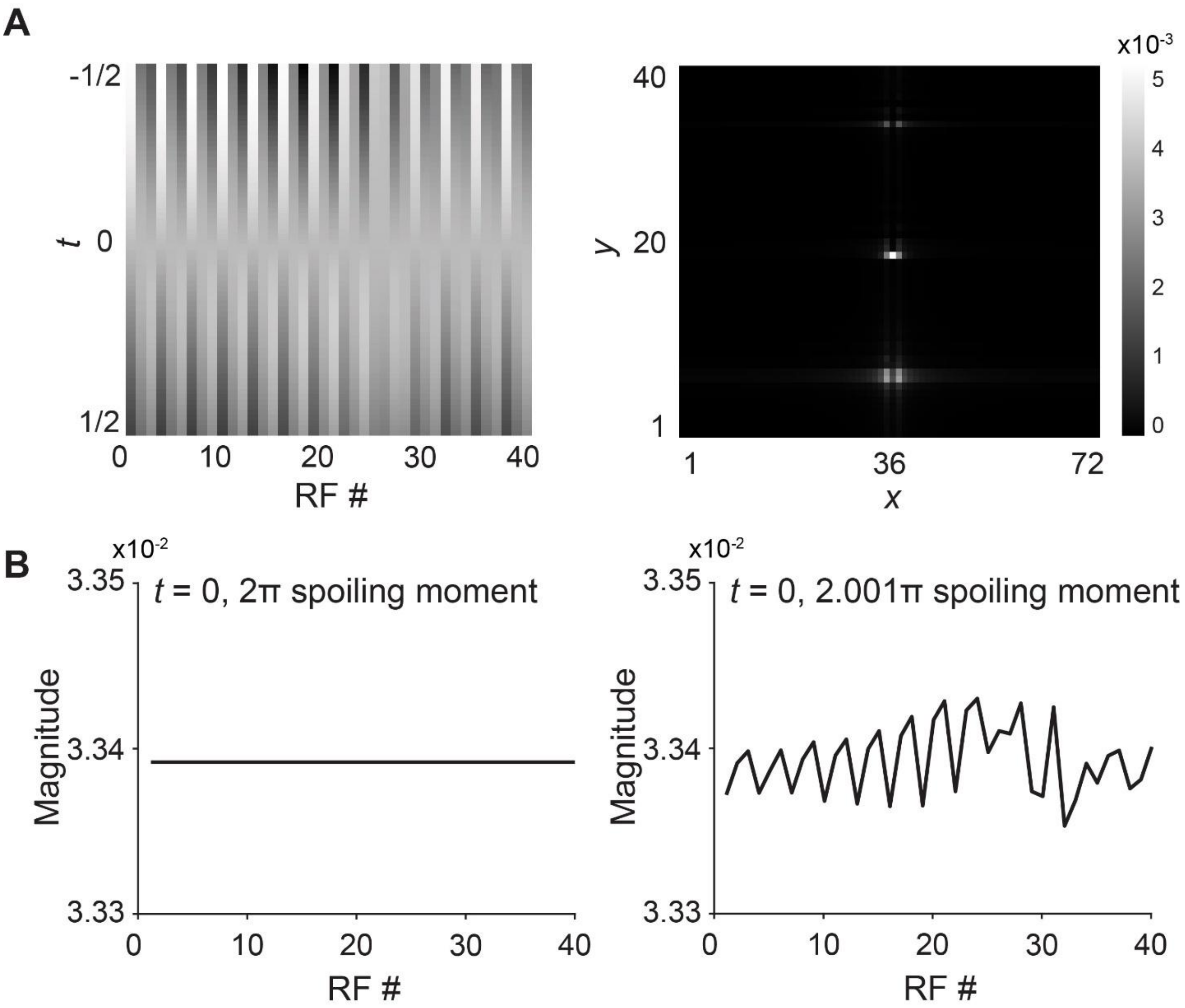


**Fig. S3.**

**Ghosting artifacts due to PSS in conventional FLASH sequence. (A)** Concatenation of 40 echoes acquired from the $2001^{st}$ to $2040^{th}$ TR by 1D Bloch simulation, to visualize signal oscillations (left). Ghosting artifacts along the phase-encoding direction in the simulated FLASH image (right). **(B)** Steady-state transverse magnetization at TE when using 2π gradient spoiling (left) and signal oscillations at TE when using 2.001π gradient spoiling (right).

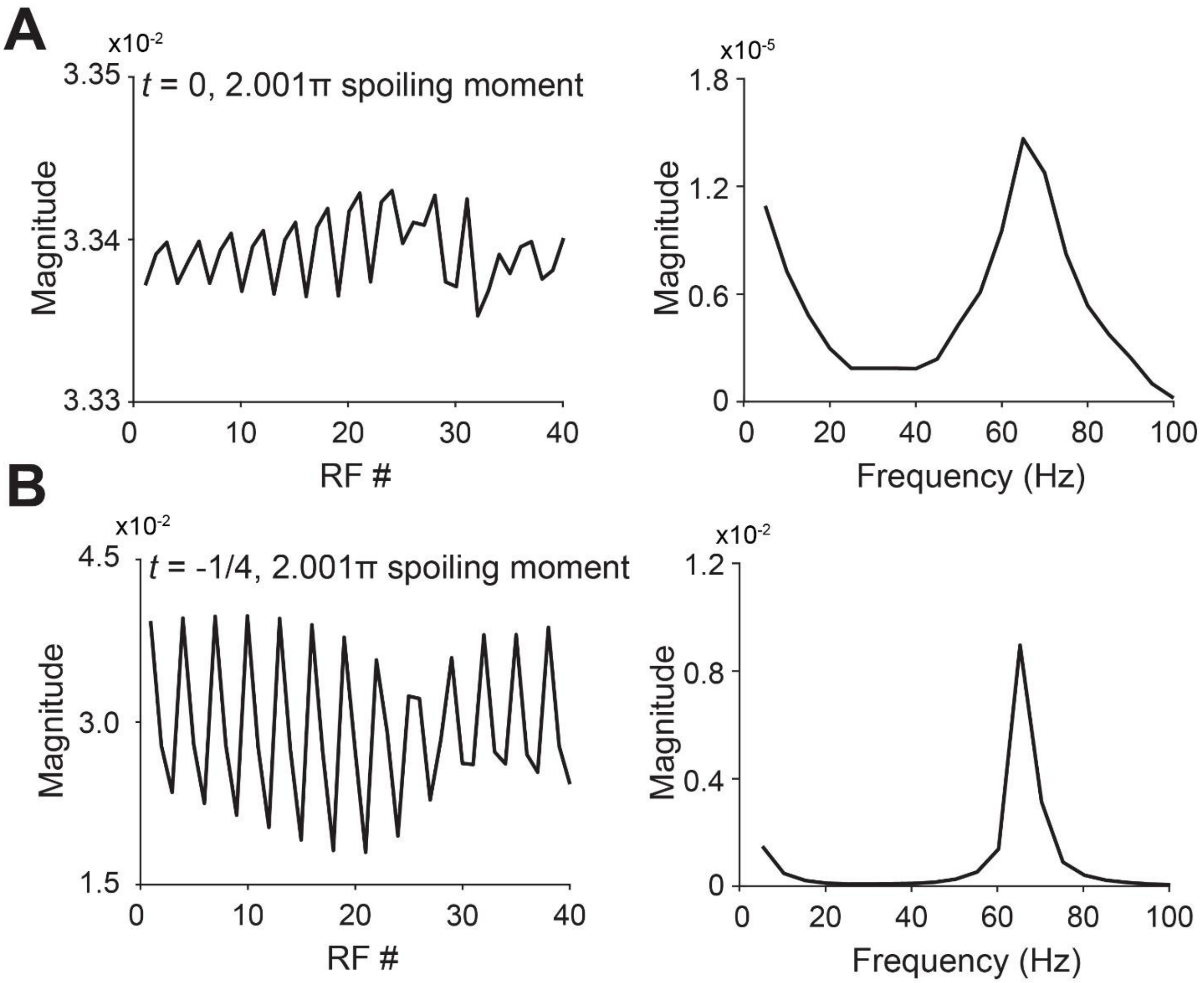


**Fig. S4.**

**PSS oscillations observed in 1D Bloch simulation and its frequency spectrum. (A)** Time series and its frequency spectrum at TE ($t = 0$) when using 2.001π gradient spoiling. **(B)** Time series and its frequency spectrum near the edges of k-space ($t = -1/4$) when using 2.001π gradient spoiling.

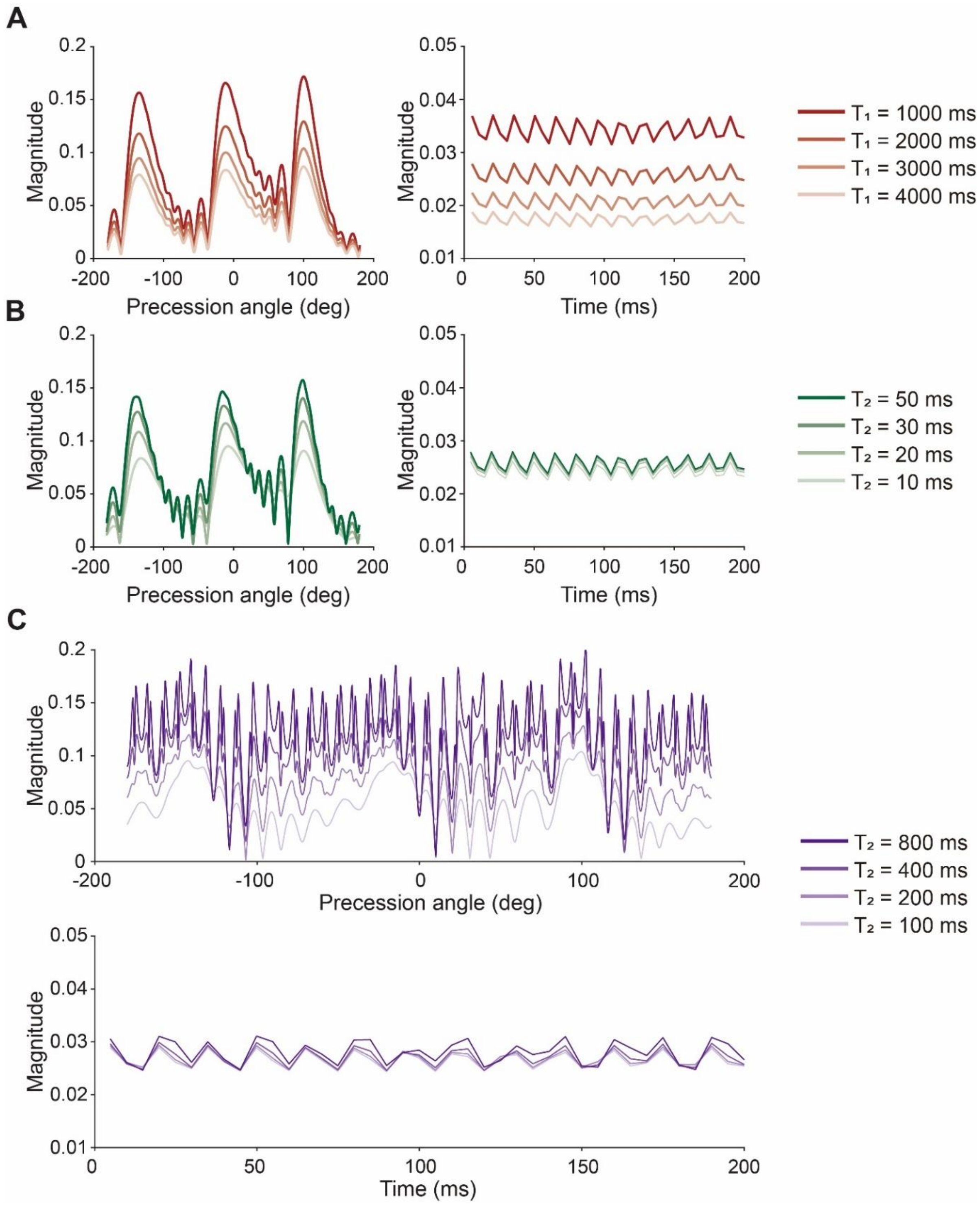


**Fig. S5.**

**Effect of $T_1$ and $T_2$ on PSS oscillation. (A)** Dependence of magnetization profiles and signal intensity within a single voxel on $T_1$. **(B)** Dependence of magnetization profiles and signal intensity within a single voxel on $T_2$. **(C)** Dependence of magnetization profiles and signal intensity within a single voxel on larger $T_2$.

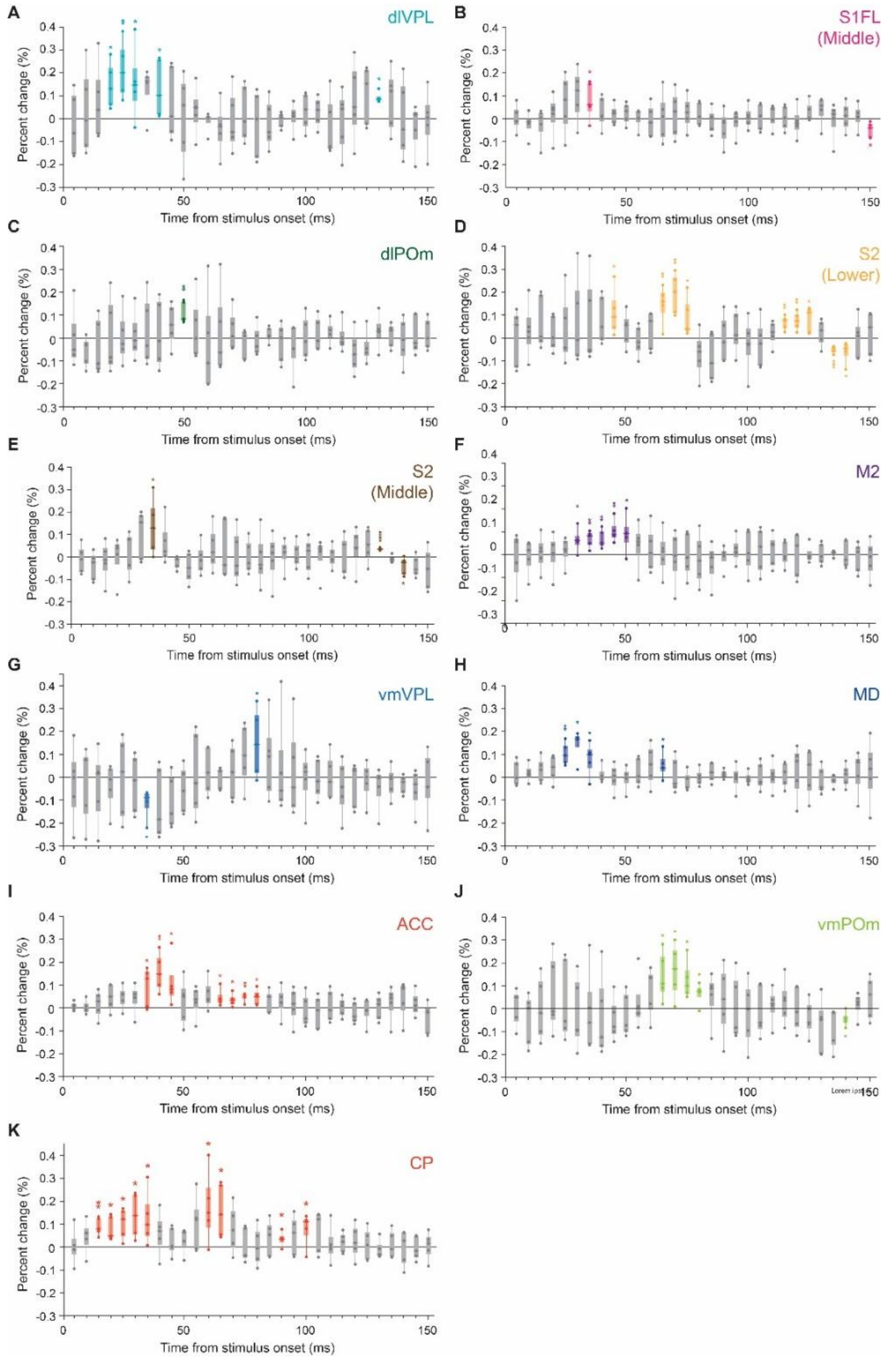


**Fig. S6.**

**Statistical significance of 11.7 T DIANA signal changes at each time point. (A to K)** DIANA responses in dorsolateral VPL (dlVPL) (A), middle S1FL (B), dorsolateral POm (dlPOm) (C), lower S2 (D), middle S2 (E), M2 (F), ventromedial VPL (vmVPL) (G), MD (H), ACC (I), ventromedial POm (vmPOm) (J), and CP (K) ($n$ = 5 mice). In the box plots, each box represents 25th to 75th percentiles, the horizontal line represents the median, and whisker represents the range from minimum to maximum. *: $p < 0.05$, **: $p < 0.01$, ***: $p < 0.001$, ****: $p < 0.0001$ for one-tailed Wilcoxon signed-rank test or one-tailed Welch's t-test.

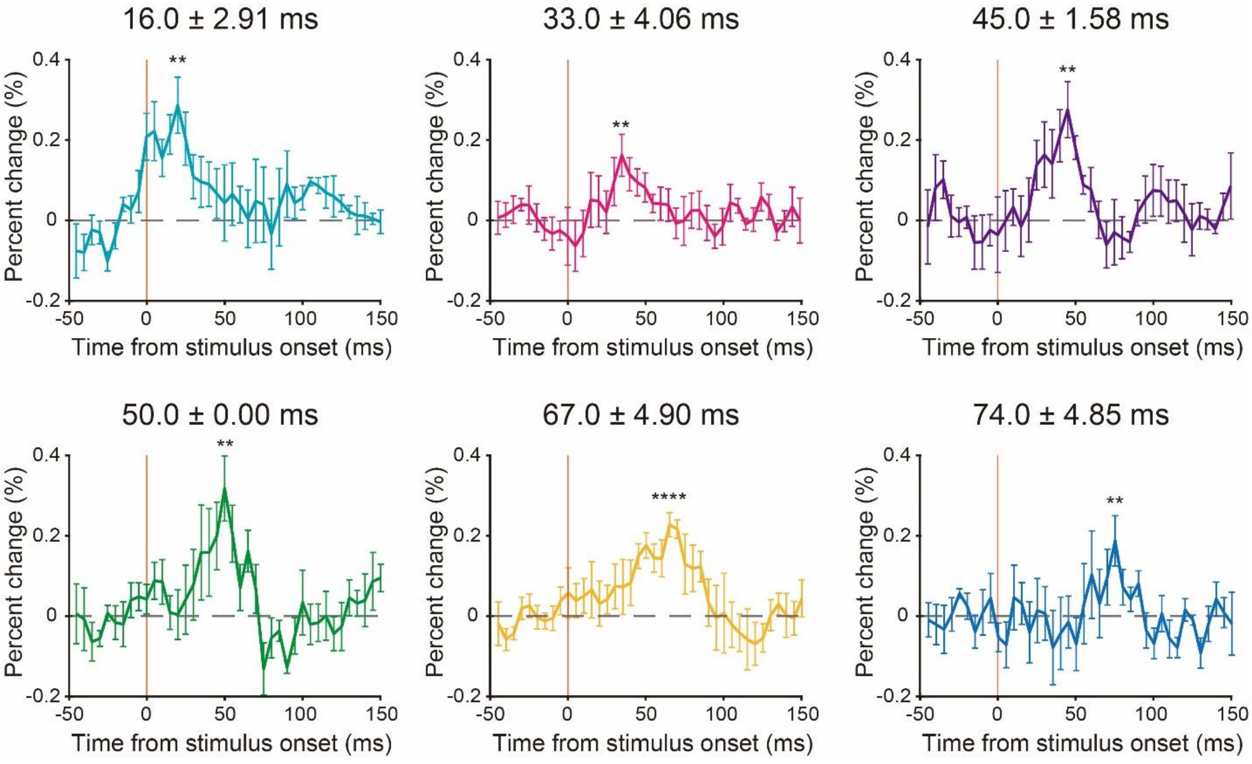


**Fig. S7.**

**Simulated DIANA responses with various peak timings.** DIANA responses were generated using 2D Bloch simulations, assigning a frequency-offset to each spin from a Gaussian distribution with a mean of 0 Hz and a standard deviation of 20 Hz ($n = 5$). Dotted red lines indicate the virtual onset time of electrical forelimb stimulation. Various peak timings similar to those *in vivo* were observed. All data are mean ± SEM. **: $p < 0.01$, ****: $p < 0.0001$ for one-tailed Welch's $t$-test.

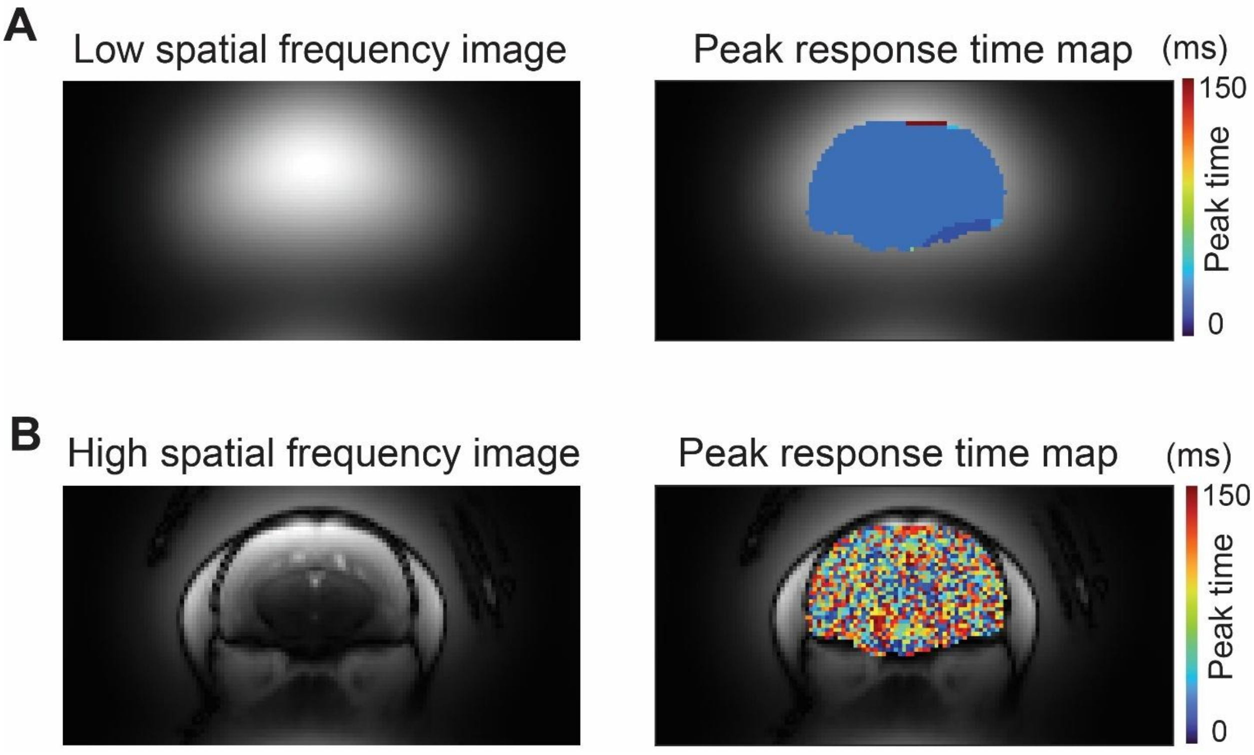


**Fig. S8.**

**Dependence of spatial frequency on the peak response time of the DIANA responses with RF spoiling and trigger delay at 11.7 T. (A)** Low spatial frequency image reconstructed using k-space around the center (left, *n* = 5 mice) and corresponding peak response time map (right). **(B)** High spatial frequency image reconstructed using the outer part of k-space (left, *n* = 5 mice) and corresponding peak response time map (right). The images consisting of only low and high spatial frequencies were reconstructed, by applying a 2D Gaussian filter centered at the center of k-space and with a FWHM of 5 and 3 data points along the $k_x$ and $k_y$ directions, respectively.
